# A cornichon protein controls polar localization of the PINA auxin transporter in *Physcomitrium patens*

**DOI:** 10.1101/2022.12.22.521699

**Authors:** C. Yáñez-Domínguez, D. Lagunas-Gómez, D.M. Torres-Cifuentes, M. Bezanilla, O. Pantoja

**Author notes:** “The author(s) responsible for distribution of materials integral to the findings presented in this article in accordance with the policy described in the Instructions for Authors is: Omar Pantoja ”. The author responsible for distribution of materials integral to the findings presented in this article in accordance with the policy described in the Instructions for Authors is: Omar Pantoj a.”.

## Abstract

Newly synthesized membrane proteins pass through the secretory pathway starting at the endoplasmic reticulum and packaged into COPII vesicles to continue to the Golgi apparatus before reaching their membrane of residence. It is known that cargo receptor proteins form part of the COPII complex and play a role in the recruitment of cargo proteins for their subsequent transport through the secretory pathway. The role of cornichon proteins is conserved from yeast to vertebrates, but it is poorly characterized in plants. To study the role of this protein in cellular traffic mechanisms in plants, the moss *Physcomitrium patens* has been selected since it can be studied at the single-cell level. Here, we studied the role of the two moss cornichon homologs in the secretory pathway. Mutant analyzes revealed that cornichon genes regulate different growth processes during the moss life cycle, by controlling auxin transport; with CNIH2 functioning as a specific cargo receptor for the auxin efflux carrier PINA, with the C-terminus of the receptor regulating the interaction and trafficking of PINA.

## Introduction

The endomembrane system of eucaryotic cells is a functionally inter-related membrane system composed of many organelles, each, with a unique membrane composition where there is a constantly exchange of proteins and lipids through a network of membrane trafficking (Morita and Shimada, 2014; Bassham et al., 2008; Jürgens, 2004; Kim and Brandizzi, 2014). Membrane trafficking comprises two main pathways: the secretory and endocytic pathways, both are essential for maintaining a wide range of fundamental cellular functions such as cell proliferation, differentiation, morphogenesis, intercellular communication and signaling, including responses to environmental stimuli (Morita and Shimada, 2014; Bassham et al., 2008). The secretory pathway involves the transport of biosynthetic materials that has been targeted to the endoplasmic reticulum (ER), then flow to the Golgi apparatus (GA) and subsequently to the plasma membrane (PM) or other organelles (Bassham et al., 2008). Newly synthesized membrane proteins in the ER are translocated to the GA by COPII vesicles (Brandizzi and Barlowe, 2013; Kim and Brandizzi, 2014). It is proposed that additional ER membrane proteins, known as cargo receptors, are required for the correct recruitment of membrane proteins to COPII vesicles, as an initial step for their transport to their target membrane. One such family of cargo receptors is the Erv14/Cornichon protein family (Dancourt and Barlowe, 2010; Herzig et al., 2012; Powers and Barlowe, 1998; Bökel et al., 2006; Castro et al., 2007; Powers and Barlowe, 2002). Cornichon (Cni) was initially identified in *Drosophila melanogaster;* during oogenesis, Cni is required for the transport of the growth factor α (TGF) Gurken (Grk) to the oocyte membrane. In the absence of *Dm*Cni, oocytes fail to establish adequate dorsoventral symmetry during oogenesis. It has been identified that Cni binds to an extracellular domain of Grk, evidence that was used to propose that cornichon acts as a cargo receptor, recruiting Grk into COPII vesicles (Bökel et al., 2006). The loss of *ERV14*, the homologue of cornichon in yeast, causes the formation of a defective budding site due to the inefficient transport of the Ax12p protein, necessary for the establishment of axial polarity (Powers and Barlowe, 1998).

CNI homologues (CNIH) proteins are also present in plants, but their role in these organisms has been scarcely studied. In rice (*Oryza sativa*) two homologous proteins, *Os*CNIH1 and *Os*CNIH2, have been identified. Using heterologous expression systems in yeast and the epidermis of tobacco leaves (*Nicotiana benthamiana*), it was observed that *Os*CNIH1 localized to the ER and GA, similar to Erv14 in yeast, and that interacts with the sodium transporter *Os*HKT1; 3, suggesting that *Os*CNIH1 functions as a cargo receptor for *Os*HKT1;3 (Rosas-Santiago et al., 2015). In *Arabidopsis thaliana*, five CNIH proteins have been identified, denominated as *At*CNIH1 to *At*CNIH5, with no obvious phenotype. In pollen, red fluorescent protein-tagged *At*CNIHs (RFP-*At*CNIHs) localized to the early secretory pathway and some of them are necessary for correct targeting of the integral membrane proteins *At*GLR2.2 and *At*GLR3.3, but not for other soluble or membrane-attached proteins, suggesting cargo specificity for pairs of *At*CNIHs (Wudick et al., 2018). From this evidence, it is proposed that CNIH function as cargo receptors for a variety of plasma membrane proteins, and in plants, they seem to play a similar role as their animal and yeast counterparts (Rosas-Santiago et al., 2015; Wudick et al., 2018).

The plant-specific family of PIN-FORMED (PIN) auxin efflux transporters are integral membrane proteins, and some of them, are polarly localized to the PM (Zazímalová et al., 2010), having an important role in regulating cell polarity processes by creating an asymmetric distribution of auxin between cells and throughout the plant (Wisniewska et al., 2006; Friml et al., 2004; Leyser, 2011; Berleth Thomas and Sachs Tsvi, 2001; Sauer et al., 2006). In *A. thaliana* it has been established that PIN polar localization and maintenance at the plasma membrane is under the control of endocytosis, polar recycling, and restriction of lateral diffusion (Dhonukshe et al., 2007; Kleine-Vehn et al., 2008). It has been demonstrated that PIN proteins are internalized via clathrin-mediated endocytosis and can be cycle back to the plasma membrane via distinct trafficking routes which involve the Trans Golgi Network (TGN) and Early Endosomes (EE) (Kitakura et al., 2011; Dhonukshe et al., 2007) mediated mainly by the Brefeldin A (BFA) sensitive-ADP Ribosylation Factor Guanine Nucleotide Exchange Factor (ARF-GEF) GNOM (Steinmann et al., 1999; Geldner et al., 2003; Naramoto et al., 2014), or by independent GNOM vias like the GNOM-LIKE1 (GNL1) (Teh and Moore, 2007), the BFA-Visualized Endocytic Trafficking Defective 1 (BEN1) (Tanaka et al., 2009), the Rab-type GTPase BEX5/RABA1B and the GEF of Rab GTPase VAN4 (Feraru et al., 2012; Naramoto et al., 2014). The retromer complex which forms a coat on the cytosolic face of endosomes, in particular the SORTING NEXIN 1 (SNX1) and VACUOLAR PROTEIN SORTING 29 (VPS29) subunits, are also required for recycling and intracellular trafficking of PINs (Jaillais et al., 2007). Recently, it was demonstrated that loss of the exocyst subunits EXO70A1 or SEC8, drastically slowed down the release of PIN from BFA compartments after drug removal, adding another mechanism in the control of PIN targeting to the plasma membrane (Drdová et al., 2013). Besides these trafficking mechanisms, little is known about the contribution of early steps of the secretory pathway that are involved specifically in the processing and sorting of de novo synthesized PIN proteins.

As CNIH proteins participate in the establishment of cell polarity through the regulation of the traffic of cargo proteins to the plasma membrane, it was of particular interest to study if trafficking of PIN proteins occurred through their interaction with the cargo receptor CNIH. Two factors have complicated functional studies of CNIH and PIN proteins in plants. First, plants have an expanded CNIH and PIN gene families and second, CNIH proteins in plants have been studied using primarily heterologous expression systems. We used the moss *Physcomitrium patens* to study CNIH-dependent cell trafficking mechanisms for several reasons. Most *P. patens* tissues are a single cell layer thick, enabling single-cell analysis within a tissue context (Reski, 1998; Cove et al., 2006; Naramoto et al., 2022). Unlike other model plants, *P. patens* presents a predominant haploid gametophytic phase and its high frequency of homologous recombination easily enables functional analysis of genes of interest (Rensing et al., 2008; Kamisugi et al., 2006). Here, we have characterized the two *CNIH* genes present in *P. patens* by generating single and double knockouts. We found that *CNIH*’s have pleiotropic effects at the gametophytic stage. We analyzed the function of moss CNIH in the early secretory pathway using green fluorescent protein-tagged proteins and confocal microscopy and demonstrated by protein-protein interaction assays that the auxin efflux transporter homologue PINA, is a cargo protein of the receptor CNIH2; in addition, we identified and characterized the role of C-terminus domains as important for the interaction and polar localization of PINA in protonema cells.

## Results

### Moss cornichon proteins are conserved and possess a longer C-terminus

The cornichon family of proteins is present in all eukaryotes, however, according to previous reports, cornichon proteins from plants and fungi are more similar than homolog proteins from animals (Rosas-Santiago et al., 2017; Nakagawa, 2019). This family of proteins play a role as cargo receptors, including in plants, as previously reported for the angiosperms rice *Os*CNIH1 (Rosas-Santiago et al., 2015) and Arabidopsis *At*CNIH1,-3 and −4 (Wudick et al., 2018). In view of this evidence, we wanted to know if cornichon proteins were also present in early diverging land plants using the bryophyte *Physcomitrium patens*. By BLAST analysis we identified two *P. patens* cornichon genes, *CNIH1* (Pp3c11_17020V3.1) and *CNIH2* (Pp3c7_11500V3.1) with homology to algae, plants, and fungi proteins (Figure 1A); each gene encodes for a protein that is 156 amino acids in length. We readily identified the IFXXL sequence motif in CNIH1 and CNIH2, which is similar to the IFRTL domain (IFX/NL in plants) (Figure 1A, black solid left bar), that serves as an interaction site with the COPII component *Sc*SEC24p in yeast (Powers and Barlowe, 2002; Pagant et al., 2015). Both moss proteins also possess the acidic domain that has been reported to be restricted to plant and fungal homologs that participates as a binding site for cargo proteins (Rosas-Santiago et al., 2017) (Figure 1A, black solid right bar). Interestingly, compared to other homologs, *P. patens* CNIH proteins possess an extended C-terminus with 15 extra amino acids characterized by the presence of several putative phosphorylation residues (Figure 1A, arrows). According to the phosphorylation prediction server NetPhos3.1 (Blom et al., 1999), in CNIH1 the three threonine residues (T146, T149, and T151) are potential phosphorylation residues (Supplemental Figure S1A) meanwhile, for CNIH2 only T149 is predicted as a potential phosphorylation site (Supplemental Figure S1B). We also analyzed the evolutionary relationship of moss CNIH with algae, plants and fungi homologs using the UPGMA algorithm (Supplemental Figure S1C). Cornichon proteins are grouped into three main categories (Figure 1B); in Group A we exclusively found cornichon homologous from chlorophyte algae which possess a lower similarity (less than 25%; shown in Supplemental Figure S1C) to their orthologs in plants and fungi; the second group corresponds to higher plants (Group P), and the third group is composed of fungal proteins (Group F). Group P is divided into four subgroups (I to IV), with moss CNIH1 and CNIH2 proteins grouping with CNIH proteins from early diverging land plant lineages such as the hepatic *Marchantia polymorpha*, and the lycophyte *Sellaginella moellendorffii* but also with *Chara braunii*, a charophyte algae (subgroup I) (Figure 1B), showing a ~50-60% similarity (Supplemental Figure S1C). Within angiosperms, CNIH proteins from monocots, like the rice *Os*CNIH2 (subgroup II), are more similar to moss CNIH’s (~40%; shown in Supplemental Figure S1C), followed by subgroup III represented by the Arabidopsis cornichon homologues *At*CNIH3, −4 and −5, and rice *Os*CNIH1. While the *At*CNIH1 and its orthologues proteins from *P. tricocarpa* (subgroup IV) are less similar to the moss cornichon proteins (~30%; shown in Supplemental Figure S1C). Together, these results indicate that in general, moss cornichons proteins are conserved as their homologues in plants, suggesting that they could play a similar function as cargo receptors in *P. patens*.

**Figure 1.**
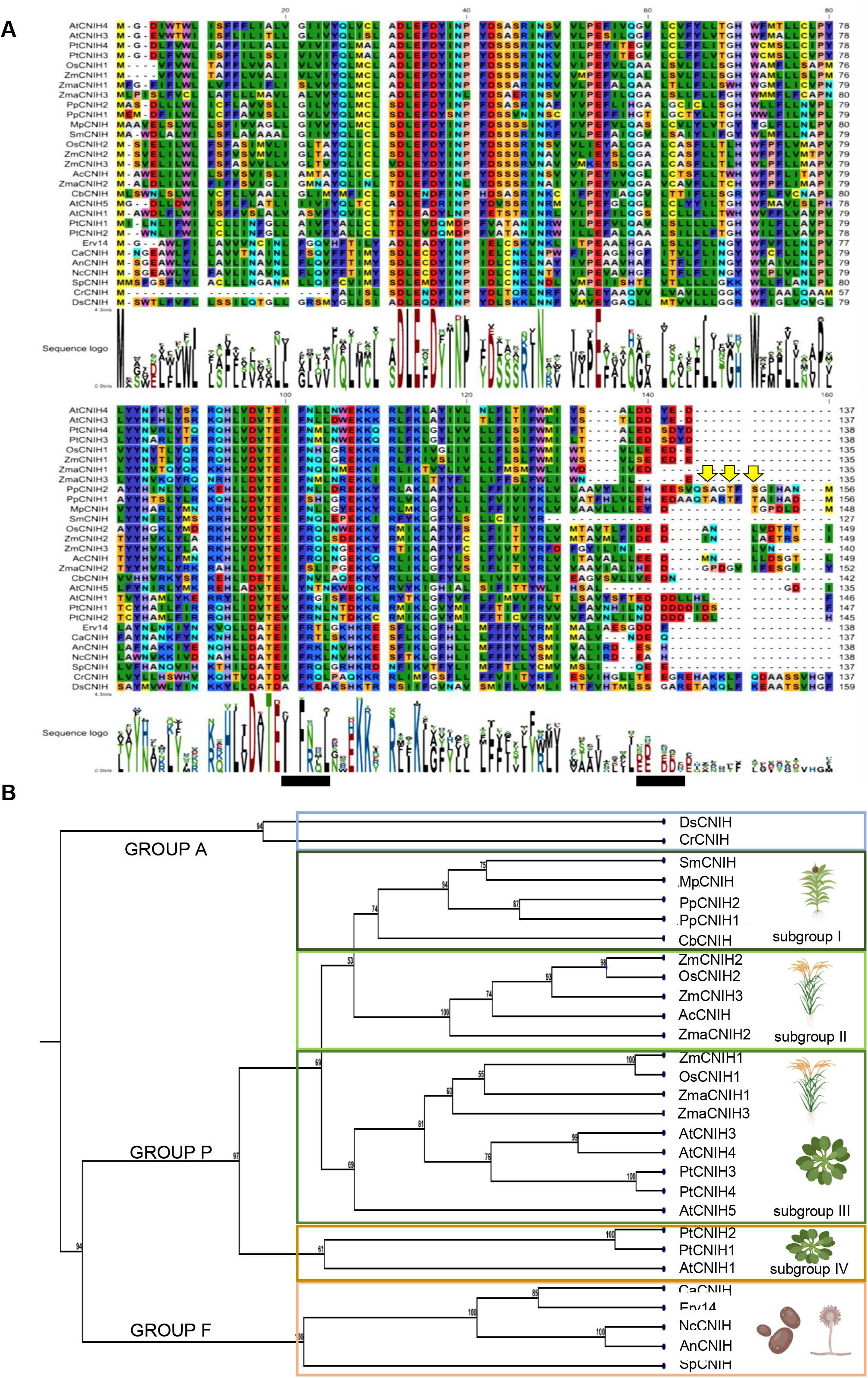
Multiple amino acid sequence alignment of cornichon homolog proteins and putative phosphorylation sites in moss homologs. **A)** Amino acid sequence alignment of cornichon homologs from algae, plants, and fungi; left and right black bars show the consensus motif IFNXL, and the acidic motif (Ac. Dom), respectively. Arrows indicate predicted phosphorylation sites on Ser and/or Thr residues in the moss homologs identified by the NetPhos3.1 prediction server. **B)** Phylogenetic tree of cornichon homologs. Group A is represented by algae, Group P, corresponds to higher plants (divided by subgroup I, grouping cornichon from bryophytes; subgroup II represented by homologs only from monocotyledons; subgroup III represented by homologs from mono-and dicotyledons; subgroup IV represented by homologs only from dicotyledons), and Group F, corresponding to fungi. The data and images were analyzed with the software CLC Main Workbench 8.1. Model organisms’ pictures are taken from Biorender templates. For nomenclature details see Materials & Methods section.

### Mutations of cornichon homologs cause subtle morphological changes along the life cycle of the moss

To understand the physiological role of the moss cornichon genes, we generated single null mutants (Δ*cnih1* and Δ*cnih2)* and a double null mutant Δ*cnih1/2* and analyzed two independent lines of each, from two independent rounds of transformation events. We used CRISPR-Cas9 system (Mallett et al., 2019) to edit *CNIH1*, resulting in an in-frame premature stop codon at nucleotide position 132, which would encode a 42 amino acid peptide (Supplemental Figure S2). For the *CNIH2*, the null mutant was generated by homologous recombination, replacing the corresponding locus by a hygromycin resistance cassette (Supplemental Figure S3). To obtain the double mutant, we used homologous recombination to replace the *CNIH2* locus with the hygromycin resistance cassette in the Δ*cnih1-23* single mutant (Supplemental Figure S2 and S3). All the mutants were viable and protonemal growth was similar to the wild type (WT) (Figure. 2A and Supplemental Figure S4A). However, we found that mutant plants exhibited abnormal branching, with side branch initials forming in the middle of the subapical cell, instead of initiating at the apical end of the subapical cell, as normally observed in WT protonemata (Figure 2A, asterisk and arrow, and Supplemental Figure S4A). To corroborate these observations, we quantified the frequency of abnormal branching, observing that approximately 13 to 18% of the branches in the *Δcnih1* and ≤10% in *Δcnih2* single mutants showed abnormal branching, while none were observed in the WT (Figure 2B and Supplemental Figure S4A) For the Δ*cnih1/2* double mutant, the phenotype was more severe, with 35 to 40% of the branching cells occurring in the middle of the subapical cell (Figure 2A-B). Additional alterations in protonemal morphology were observed in the Δ*cnih2* mutant, including an increased number of side branch cells (Figure 2C) and early development of caulonemal cells, confirmed by the larger calculated caulonemal/chloronemal cell ratio (Figure 2D). These results suggest that both *CNIH1* and *CNIH2* help to regulate positioning of the branch initial cells in protonemata; in addition, *CNIH2* seems to play a role in negatively regulating protonemal branching, and in cell differentiation from chloronemata to caulonemata.

**Figure 2.**
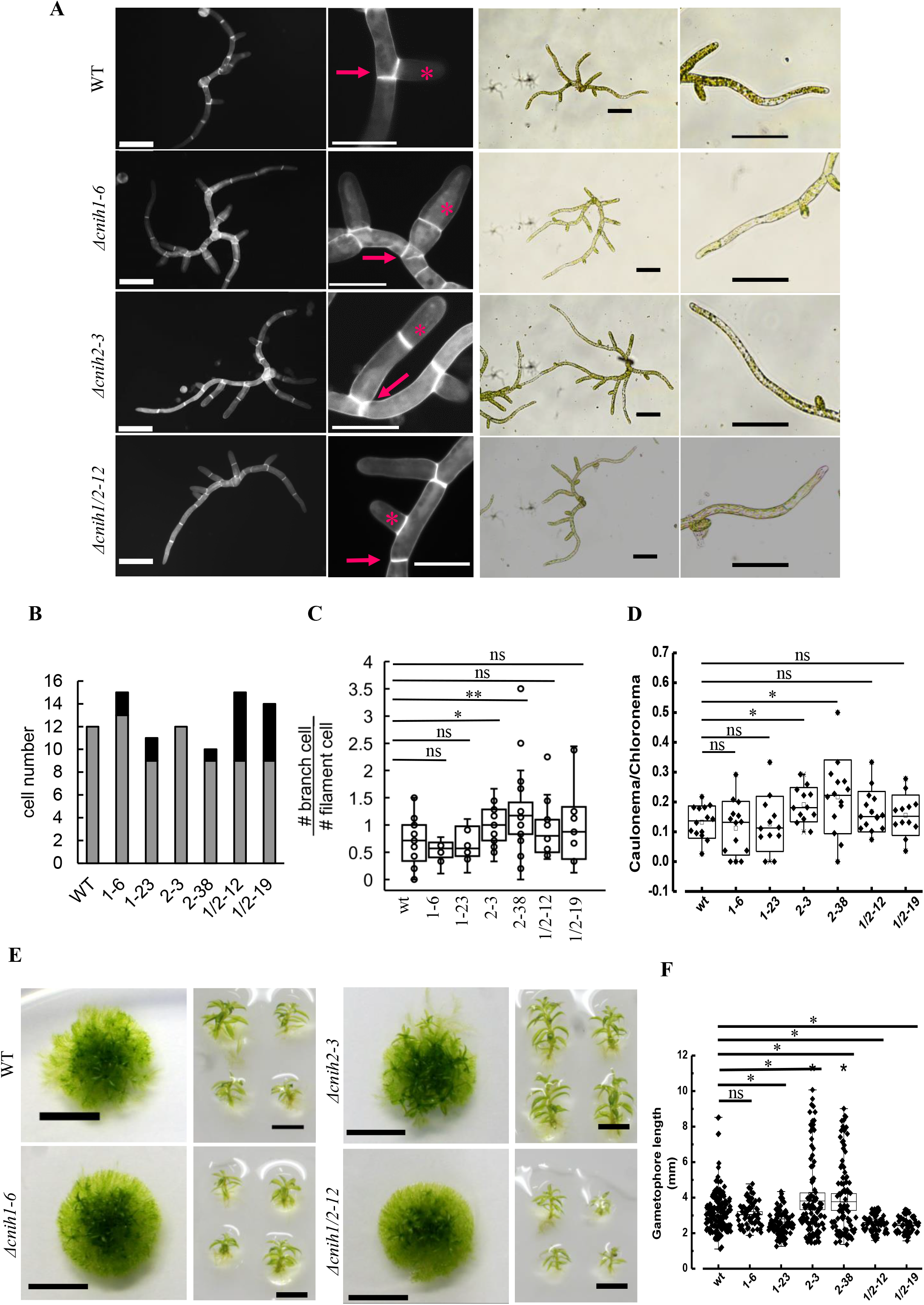
Cornichon mutants have pleiotropic effects during the gametophyte moss life cycle. **A)** (Left panel) Protonema from WT and mutant lines stained with Calcofluor White after seven-day growth, visualized under epifluorescence microscopy; scale 100 μm. Enlargements show cell division (arrow) and lateral initial branch cell (*). (Right panel) Brightfield images from seven-day-old protonema and enlarged protonema tip zones; scale 50 μm. **B)** frequency of lateral cells emerging at the middle of the subapical protonema cell by plant at 7 d, n ≥ 10; gray bars indicate frequency of normal lateral cells and black bars indicate frequency of anormal lateral cells. **C)** Branching ratio of protonema lines, by quantifying the number of total lateral cells divided by total cell number per filament at 7 d, n ≥ 11. **D)** Calculated caulonema/chloronema ratio, n > 12; *t*-test was performed for protonema statistics (ns p ≥ 0.05; * p < 0.05; ** p ≤ 0.001). **E)** Colony (left; scale 5 mm) and individual gametophores (right; scale 2 mm) from WT and mutant lines at four weeks of growth. **F)** Gametophore size of the lines, n ≥ 59. ANOVA and Tukey-Kramer post hoc test was performed for statistics (ns p ≥ 0.05; * p < 0.05).

Gametophores from the *Δcnih2* mutant were bigger than WT, while those from the Δ*cnih1* and Δ*cnih1/2* mutants were smaller than the WT (Figure 2E-F and Supplemental Figure S4B). Among the phenotypes observed from the three cornichon mutants, the early caulonemal development in the *Δcnih2* mutant was of particular interest because it has been suggested that polar transport of auxins mediated by auxin efflux transporters (PIN) are important for the transition from chloronemata to caulonemata (Viaene et al., 2014; Bennett et al., 2014). Interestingly, deletion of the PINB transporter also generated larger gametophores like the Δ*cnih2* mutant (Bennett et al., 2014). These similarities led us to explore a possible interplay between moss CNIHs and the PIN transporters.

### CNIH2 is the cargo receptor for the auxin efflux transporter PINA that controls protonemal development

To investigate a possible link between cornichon and the PIN transporters, we analyzed protein-protein interactions between the cornichon homologs and one of the auxin transporters, PINA, the main isoform expressed in all moss tissues (Bennett et al., 2014). Since PINs and cornichons are integral membrane proteins, we employed the mating-based Split Ubiquitin System (mbSUS) that is designed to identify interactions between membrane proteins (Obrdlik et al., 2004; Lalonde et al., 2010). We found that CNIH2 interacted more strongly with PINA than CNIH1, as indicated by enhanced growth on selection medium (Figure 3A, IS-0) and lower inhibition caused by Met (Figure 3A, IS-500). We also observed greater LacZ activity for the interaction between CNIH2 and PINA as compared to CNIH1 and PINA (Figure 3A, LacZ). Additionally, by employing Bimolecular Fluorescence Complementation (BiFC) in *Nicotiana benthamiana* epidermal cells, we confirmed that CNIH1 and CNIH2 interacted with PINA on reticulated structures that resemble the ER, as well as puncta distributed throughout the cytoplasm (Figure 3B). The well-established homo-oligomerization of the aquaporin *At*PIP2 (Christophe Maurel et al., 2015), was used as a positive control for the BiFC assay (Figure 3B). The interaction between cornichon and the aquaporin agreed with similar interactions identified in Arabidopsis (Jones et al., 2014). Coexpression of the auxin transporter and the aquaporin resulted in no observable fluorescence, indicating that PINs do not interact with aquaporins and served as a negative control (Figure 3B). Given the interaction between cornichon and the auxin transporters, we hypothesized that auxin transporter trafficking might be altered upon cornichon mutation, the putative receptors. To test this, we mutated *CNIH1* or *CNIH2* using CRISPR-Cas9 or homologous recombination, respectively, in the moss reporter line PINA-EGFP (Viaene et al., 2014). We obtained two independent lines for each from two different transformations (Supplemental Figure S2-3). Without *CNIH1*, PINA-EGFP localization was not modified (Figure 3C and Supplemental Figure S5A); however, deletion of *CNIH2* resulted in mislocalization of the auxin transporter, as indicated by the absence of the apical fluorescence associated with PINA-EGFP (Figure 3C and Supplemental Figure S5A). To confirm that apical PINA depended on CNIH2, we transiently expressed the full coding sequence of *CNIH2* with the ubiquitin promoter in the *Δcnih2*/PINA-EGFP mutant line and observed restoration of PINA apical localization (Figure 3C). These results demonstrate that CNIH2 is required for PINA trafficking to the plasma membrane in protonemal cells.

**Figure 3.**
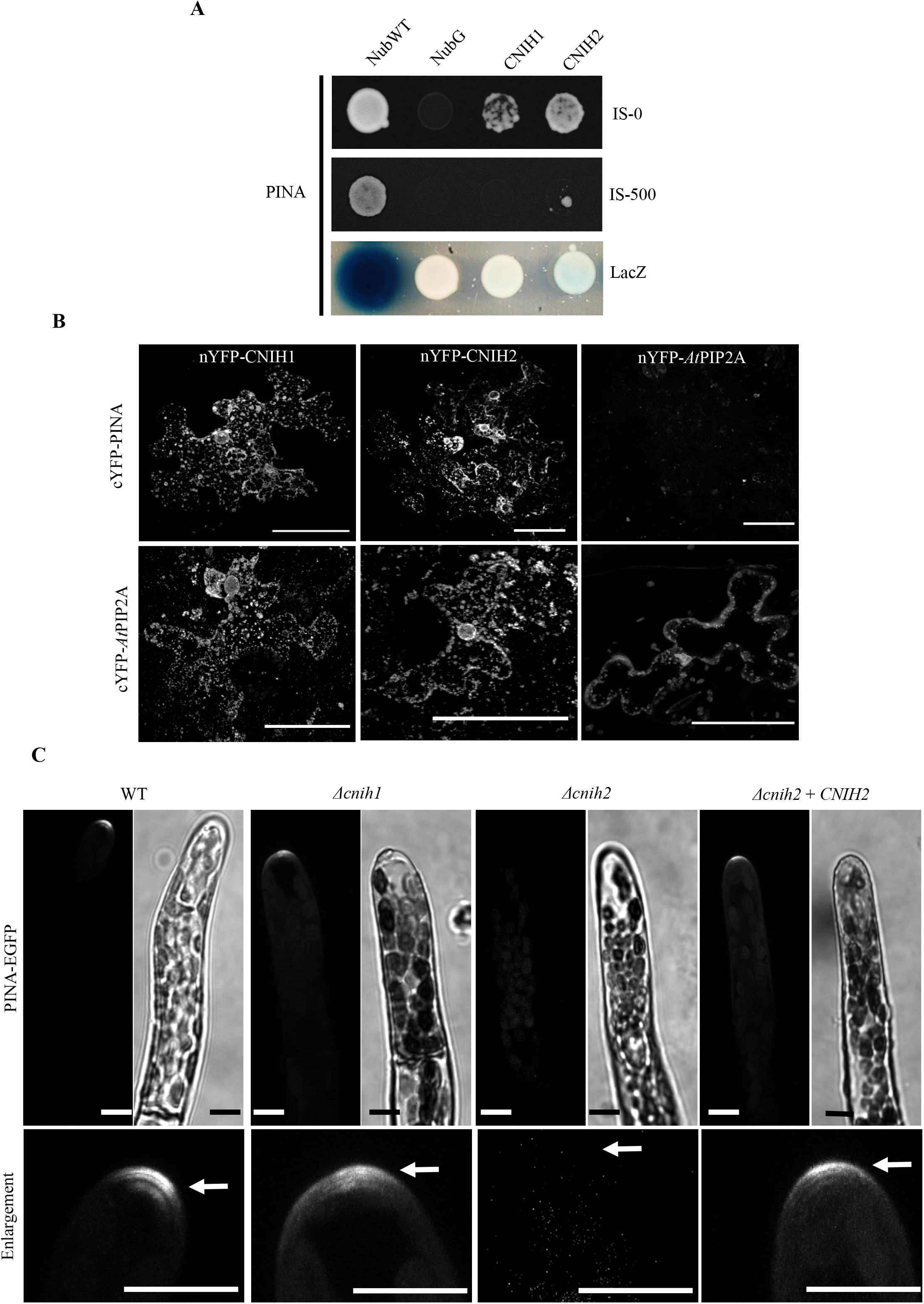
CNIH2 protein is the cargo receptor for the auxin efflux carrier PINA. **A)** Protein-protein interaction identified by the mbSUS assay with the moss cornichon proteins (Nub fusions) and the auxin transporter PINA (Cub fusion). Yeast cell growth in selection medium (IS-0); the differential interaction was confirmed by cell growth inhibition under repressive selection conditions (0.5 mM Met; IS-500) and by the lower activity of LacZ (intensity of the bluish color). NubWT and NubG were used as false negative and false positive controls, respectively. **B)** Interaction between PINA and CNIH1 or CNIH2 was confirmed by reconstitution of split-YFP fluorescence by the co-expression of nYFP-CNIH’s with c-YFP-PINA protein. The lack of interaction between cYFP-PINA and the aquaporin nYFP-*At*PIP2A was indicated by the absence of fluorescence (negative control). Oligomerization of the aquaporin (nYFP-*At*PIP2A and cYFP-*At*PIP2A) was used as positive control. The interaction between the aquaporin cYFP-*At*PIP2A and nYFP-CNIH’s was evinced by the reconstitution of YFP fluorescence. Scale = 50 μm. **C)** Apical localization of PINA-GFP in WT was unaffected in the *Δcnih1* mutant but delocalized in the *Δcnih2* mutant, a phenotype recovered by complementation with *CNIH2*. Bottom panels are ROI enlargements of apical protonema cells from the corresponding GFP images; arrows indicate PINA-GFP localization. Images are z-projections with the maximal intensity. Scale 10 μm.

### CNIH2-associated puncta are insensitive to Brefeldin A and are associated with a SEC23G subpopulation of ER exit sites (ERES)

The cornichon family of proteins has been characterized as cargo receptors in the ER for the selection of cargo membrane proteins to be transported to the Golgi as part of the early secretory pathway (Herzig et al., 2012; Castro et al., 2007; Rosas-Santiago et al., 2015). To determine the localization of CNIH2 in moss cells, we used CRISPR-Cas9 in combination with homologous directed repair (CRISPR-Cas9 & HDR) (Mallett et al., 2019) to insert three tandem sequences encoding for mRuby2 (hereafter, 3XmRuby) in-frame with the coding sequence of *CNIH2* at the endogenous locus (Supplemental Figure S6 A, C-D). We found that CNIH2-3XmRuby localized to puncta throughout the cell and at the ER, we also observed the puncta localized around the nucleus, which corresponds to perinuclear ER, and a particular accumulation of the puncta near the apex of protonemal cells (Figure 4A, right). CNIH2-3XmRuby was also localized at the cell periphery in protonemal cells, corresponding to cortical ER (Figure 4A, left, arrows). This subcellular localization was consistent with what we observed in protonemal cells transiently overexpressing the *ZmUBIpro:*CNIH2-EGFP or *ZmUBIpro:*CNIH1-EGFP fusions (Supplemental Figure S7). To identify the organelle associated with the CNIH2-3XmRuby puncta, we treated protonemal cells with Brefeldin A (BFA), a drug that disassembles the Golgi and causes its redistribution back into the ER (Roberts et al., 2018; Ito et al., 2012; Chardin and McCormick, 1999). Employing the cis-Golgi marker line (YFP-GmMan1), we confirmed that Golgi-associated vesicles were disrupted by BFA (Figure 4B, right panels). Under this condition, however, CNIH2-3xmRuby puncta were still observed (Figure 4B, left panels), indicating that they do not correspond/localize to the GA. It is well established that cargo proteins are loaded into ER subdomains known as ER exit sites (ERES), where the COPII subunits Sec23, Sec24, Sec13 and Sec31 are recruited (Brandizzi and Barlowe, 2013). Recently, three protonemal-expressed Sec23 isoforms were shown to associate with the ER, with SEC23G forming larger puncta as compared to SEC23B and SEC23D (Chang et al., 2021). The SEC23G puncta were remarkably similar in size to CNIH2-associated puncta. To test for colocalization of CNIH2 and SEC23G in protonemal cells, we employed CRISPR-Cas9 & HDR to insert three tandem sequences encoding for mNeon (hereafter, 3XmNeon) in-frame with the coding sequence of *SEC23G* at the endogenous locus in the *CNIH2-3xmRuby* line (Supplemental Figure S6 B-D). Using confocal microscopy, we observed partial colocalization of CNIH2-3XmRuby with SEC23G-3XmNeon (Figure 5A), as indicated by the calculated Pearson’s correlation coefficient of 0.66 (Figure 5B). To corroborate that the co-localization analysis was significant, we flipped horizontally or vertically the Sec23G images and repeated the co-localization analysis and found that the correlation coefficients were significantly lower (Figure 5B). Coincidentally, the puncta individually highlighted by both proteins shared a similar size, but smaller than those associated with the Golgi (Figure 5C). These results suggest that the puncta associated to CNIH2 may correspond to SEC23G-structured ERES and that the polarized localization of moss PINA is mediated by the early secretory pathway through its association with CNIH2.

**Figure 4.**
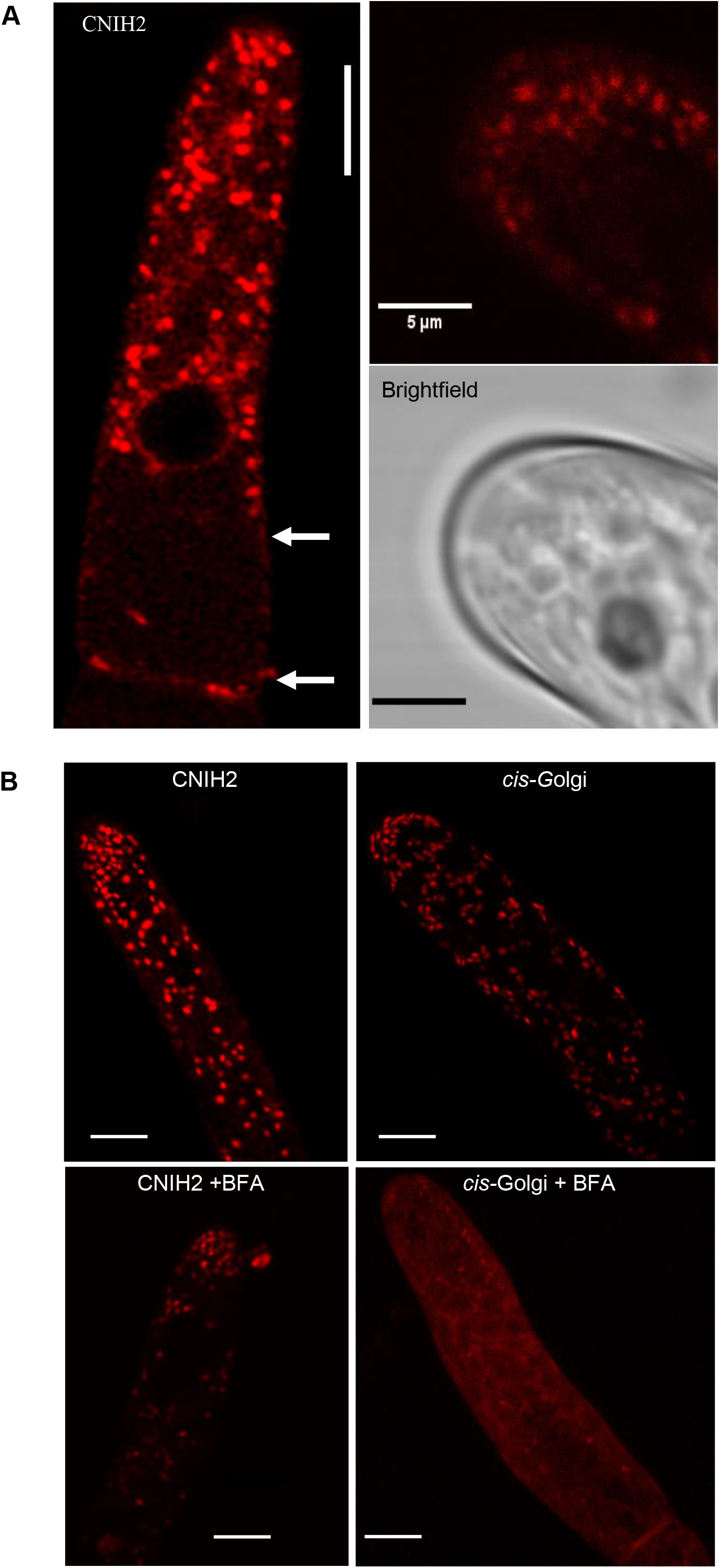
CNIH2 localizes at the ER and puncta insensitive to BFA. **A)** Subcellular localization of endogenous CNIH2 at the endoplasmic reticulum and associated puncta in the apical protonemal cell (left) and concentrated at the apex zone (right); confocal image taken from the *CNIH2-3xmRuby* transgenic moss line; arrows indicate peripheral localization at ER; scale 10 μm. **B)** Subcellular localization of CNIH2-3xmRuby (left) and cis-Golgi (YFP-GmMan, right) in an apical protonemal cell before (top) and after exposure to 50 μM Brefeldin A for 24 h (+BFA; bottom); scale 10 μm.

**Figure 5.**
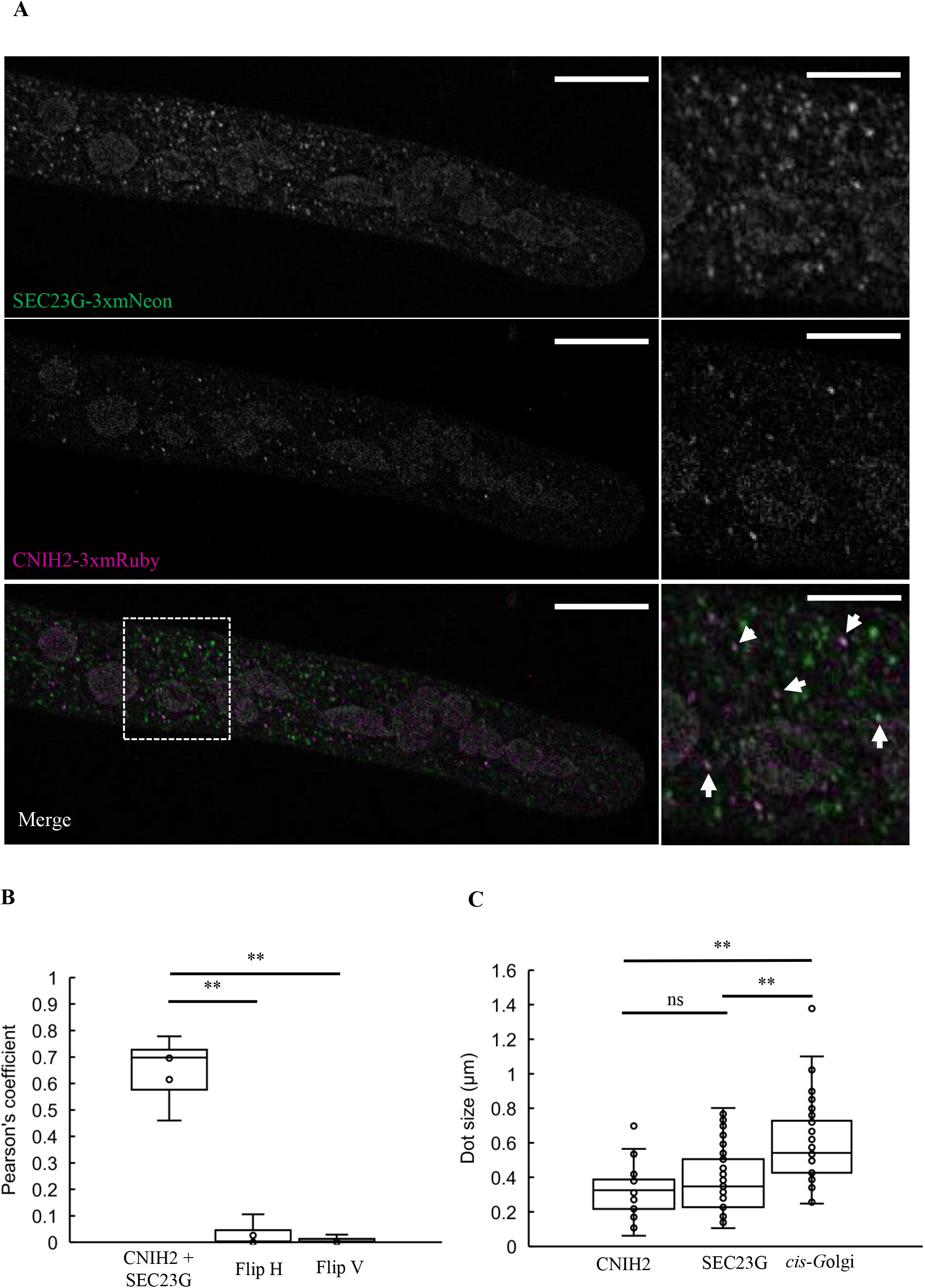
CNIH2 localizes at SEC23G confined ERES. **A)** (left) Co-localization of endogenous CNIH2 and SEC23G tagged proteins in a protonemal apical cell; (right) enlargement of the region delimited (dashed square), arrows show co-localization of both proteins. Representative Z-projection with maximal intensity confocal image; scale 10 μm. **B)** Pearson’s correlation coefficient measured from endogenous CNIH2 and SEC23G tagged proteins in comparison with horizontal (Flip H) and vertical flipped (Flip V) SEC23G images. *t*-test was performed for statistics (p ≤ 0.001); n = 6 cells. **C)** Vesicle size for CNIH2 (0.38 ± 0.18 μm), SEC23G (0.32 ± 0.13 μm) and cis-Golgi (0.59 ± 0.24 μm) from protonemal cells; n = 35. Data are the mean ± SD; *t*-test was performed for statistics (ns, p ≥ 0.05; *, p ≤ 0.05).

### The C-terminus of CNIH2 regulates its interaction with PINA and trafficking of the transporter

Correct trafficking of cargo membrane proteins depends on specific domains present in the Erv14/CNIH protein family (Pagant et al., 2015; Rosas-Santiago et al., 2017), which led us to analyze domains in CNIH2 that might influence PINA trafficking. One of the characteristics of moss CNIH homologs is the presence of a long C-terminus with an acidic domain, and a domain containing potential phosphorylation sites (Figure 1). To investigate if these domains regulated CNIH2 function, we generated two truncations, one removing the domain containing the putative phosphorylation sites, denoted T/S, at the extreme C-terminus (CNIH2-141), and the second, removing the T/S and the acidic domain (CNIH2-137) (Figure 6A). To determine if the truncations affected CNIH2 interactions with PINA, we employed the mbSUS assay. As shown in figure 6B, the strength of the interaction was enhanced when the T/S domain was removed (CNIH2-141), as indicated by cell growth in the presence of 0.5 mM Met but reduced when both domains were removed (CNIH2-137). These results suggest that the T/S domain is involved in regulating the strength or the stability of the interaction with the cargo, however, it remains to be demonstrated if the phosphorylation state of T149 (Supplemental Figure S1B) is indeed involved in this response. These results also demonstrated that the conserved acidic domain at the C-terminus is important to maintain the interaction with the cargo, which agrees with previous results from plants and fungi (Rosas-Santiago et al., 2017). Even with a complete C-terminal deletion, the receptor maintained a weak interaction with the cargo (Figure 6B; 0 μM Met), indicating the possible participation of other interaction sites that have yet to be identified. To corroborate if the CNIH2 truncations were expressed properly in moss, we transiently expressed the WT and C-terminus truncations of CNIH2 fused to GFP in moss protoplasts and found that WT CNIH2 and the respective C-terminus truncations were expressed at the ER and in puncta (Supplemental Figure S8). Based on these results and to ascertain the physiological importance of the C-terminus of CNIH2, we investigated if the truncations influenced the localization of PINA by transiently transforming and expressing the coding sequence of WT *CNIH2* or the truncated coding sequences of *CNIH2* driven by the maize ubiquitin promoter in the mutant *Δcnih2*/PINA-EGFP line. After maintaining plants for twelve days on selection, the protonemata from surviving plants were analyzed. The characteristic polarized localization of PINA-EGFP restricted to the tip of the apical cell was reconstituted by transformation with WT *CNIH2* (Figure 7A, left); however, transformation with *CNIH2-141* resulted in diffuse PINA-EGFP fluorescence that covered a larger area of the tip, and was less restricted to the apex of the apical cell (Figure 7A, middle). Transformation with *CNIH2-137* resulted in even more diffuse PINA-EGFP fluorescence, which was distributed all over the apical cell, labelling a structure that resembled the ER and surrounding the nucleus, suggesting that PINA was retained in the ER (Figure 7A, right, asterisks). To quantify and confirm the intracellular distribution of PINA-EGFP, a region of interest (ROI) covering the tip of the apical cell was selected to obtain a fluorescence intensity histogram (Figure 7B). In the apical cell expressing the full-length *CNIH2*, PINA fluorescence peaked at lower values (low or no fluorescence), indicating a limited localization of PINA, corresponding to the cell apex, in contrast, *CNIH2-141* and *CNIH2-137* exhibited broader peaks of PINA-EGFP fluorescence, indicating a wider intracellular distribution of PINA (Figure 7B). Together, these results suggest that the putative phosphorylation and acidic domains of CNIH2 are important for the interaction with the cargo and correct trafficking of the auxin transporter to the apical plasma membrane. Interestingly, the protonemal growth was altered in plants expressing the C-terminal truncations. In contrast to straight protonemata observed when WT *CNIH2* was expressed, expression of the truncations produced progressively curvier protonemata, with *CNIH2-141* producing wavy filaments (Figure 7C, arrow) and CNIH2-137 producing zigzagging filaments (Figure 7C, arrow). It is likely that mislocalization of the PINA auxin transporter results in changes in the direction of auxin efflux, inducing an undulatory growth in protonemata. This effect is reminiscent of the rapid changes in polar localization of Arabidopsis PIN proteins in response to environmental or developmental cues such as embryonic development or root gravitropism (Kleine-Vehn et al., 2010).

**Figure 6.**
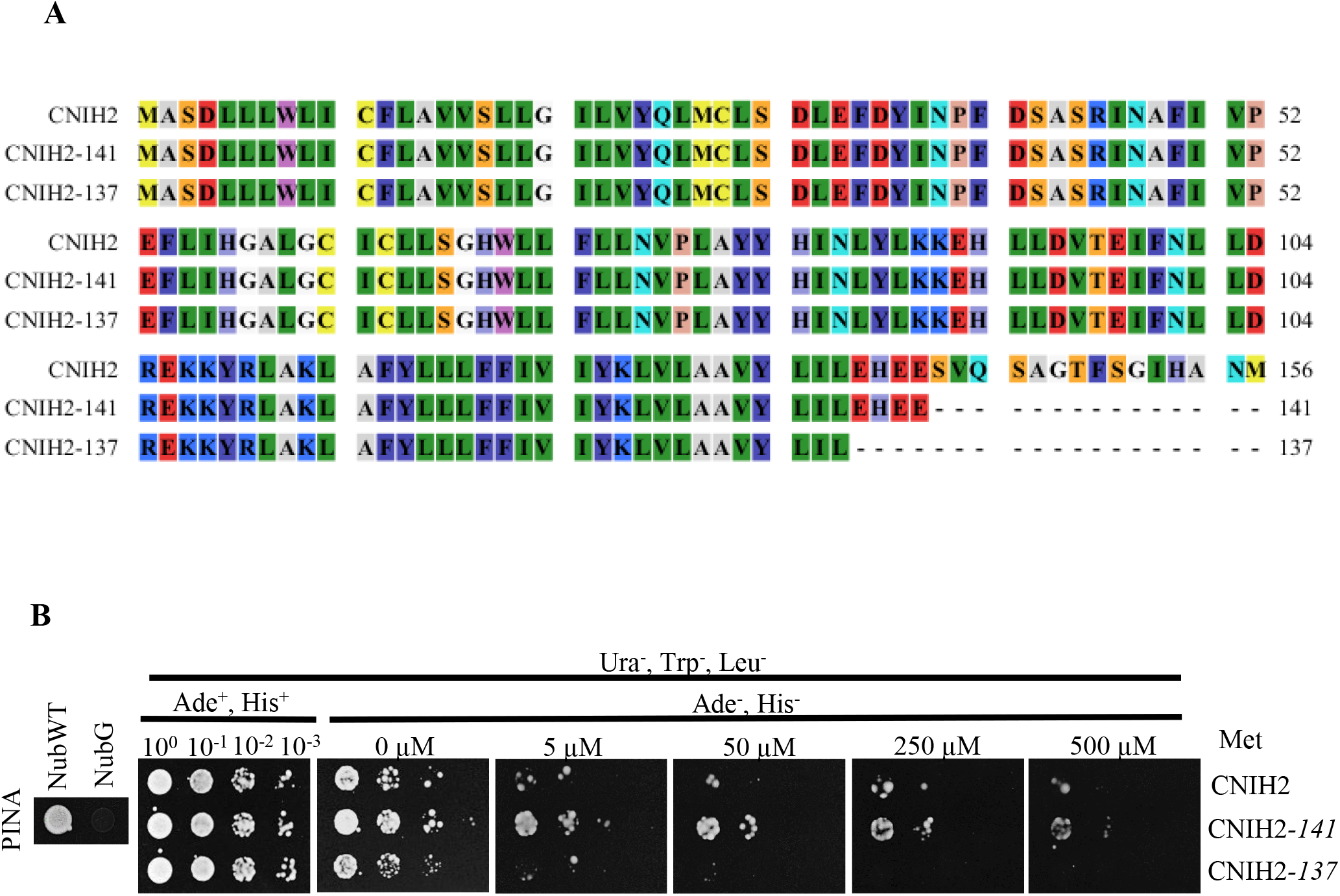
The C-terminus of CNIH2 is important for the protein-protein interaction with PINA. **A)** Amino acid sequences for CNIH2 and truncated C-terminus proteins; CNIH2-141 (T/S domain removed) and CNIH2-137 (T/S and acidic domains removed). Image generated with the software CLC Main Workbench 8.1. **B)** mbSUS assay indicated the enhanced interaction of the cargo PINA (Cub fusion) with CNIH2-141 (Nub fusion) and a diminished interaction with CNIH2-137 (Nub fusion) under increasing concentrations of Met, in comparison with CNIH2. NubWT and NubG were used as false negative and false positive controls, respectively. Growth in the presence of Ade and His corresponds to control growth conditions.

**Figure 7.**
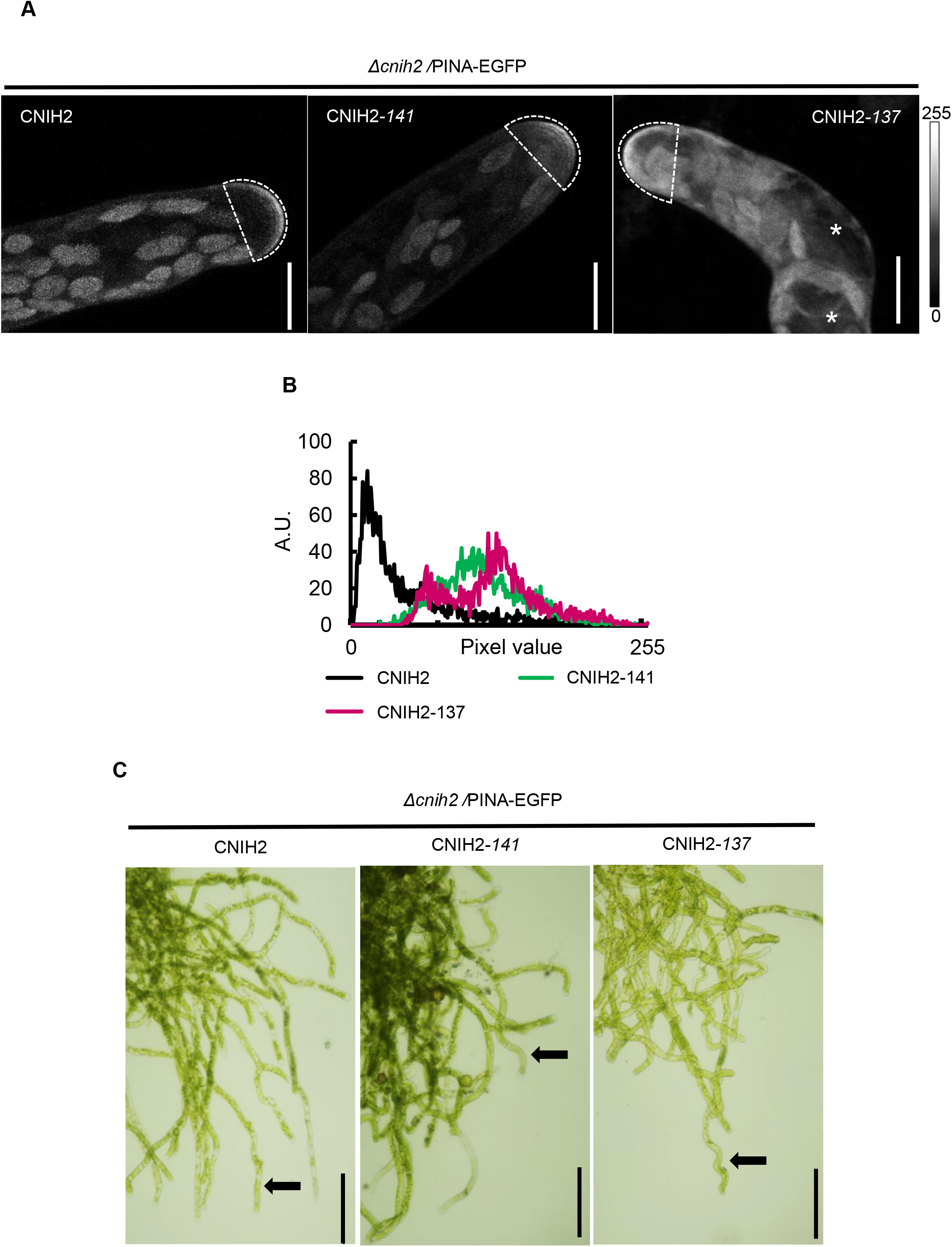
The C-terminus of CNIH2 regulates PINA trafficking in protonema cells. **A)** Changes of PINA-GFP localization by complementing with the *CNIH2* full-length and truncated *CNIH2-141* and *CNIH2-137*, using the mutant *Δcnih2-3*/PINA-EGFP moss reporter line as genetic background. Confocal images of protonema apical cells show changes in the polar localization of PINA. Images are Z-projections with maximal intensity. Asterisks shows PINA-GFP at ER localization; Scale 10 μm. **B)** Quantification of fluorescence intensity from a ROI delimiting the apex of the cells. Pixel values 0 to 255 in a gray 8-bit scale. **C)** Complementation of mutant *Δcnih2-3*/PINA-EGFP plants with the full-length CNIH2 showed normal growth, however, when complemented with truncated *CNIH2-141* or *CNIH2-137* truncated coding sequences generated an undulating protonema (arrows). Scale 100 μm.

## Discussion

Here we demonstrate that the polarized localization of PINA depends on its interaction with the cargo receptor CNIH2 via the secretory pathway (Figures 3, 4 and 5). We show that among the two cornichon homologs present in *P. patens*, CNIH2 establishes a stronger interaction with PINA (Figure 3A), an observation that correlates with PINA mislocalization in plants lacking CNIH2 but not of its paralog, CNIH1 (Figure 3C and Supplemental Figure S5). Moreover, we demonstrate that the C-terminus of CNIH2 seems to play an important role in its interaction with the cargo protein PINA (Figure 6), as deletion of a domain containing putative phosphorylation sites (CNIH-141) leads to stronger interactions and partial retention of the auxin transporter in the ER (Figure 7A-B); additionally, removal of the acidic domain (CNIH-137) exacerbates this effect by completely preventing the apical localization of PINA in the apical cell of the protonemata causing a clear retention of the transporter in the ER (Figure 7A-B). Importantly, confirmation that the correct localization of PINA is dependent on its interaction with CNIH2 was obtained by the growth defects that were caused by the truncated versions of the cargo receptor, leading to undulating growth of protonemata when expressing either truncation (Figure 7C).

A key point in the secretory pathway is the endoplasmic reticulum, where the exit of membrane proteins is under the control of the COPII coat or directly by cargo receptors (Brandizzi and Barlowe, 2013; Barlowe and Miller, 2013). In this report we identified that moss cornichons are located at the ER and in puncta in protonemata cells (Figure 4A and Supplemental Figure S7A); we demonstrated that the moss auxin efflux transporter PINA is a cargo protein for the moss cornichon cargo receptors, CNIH1 and CNIH2, interactions that occurred at the endoplasmic ER and in puncta overall the cytoplasm (Figure 3B), suggesting that moss cornichon proteins have a conserved role in early secretory trafficking. These observations are in agreement with those reported for the cargo receptor Erv14p in yeast, which shows a predominant ER localization where it interacts with cargo membrane proteins for their recruitment to the ERES through its interaction with Sec24p and eventually, within the COPII vesicles, before reaching the Golgi (Powers and Barlowe, 1998, 2002; Herzig et al., 2012; Pagant et al., 2015), reinforcing our conclusion that *Pp*CNIH2 is the preferential cargo receptor for the auxin transporter PINA. Additional evidence supporting this conclusion is the higher affinity of PINA for CNIH2 (Figure 3A), which could explain the alterations in early caulonemal development observed in the *Δcnih2* mutant (Figure 2A, D), as a result of PINA mislocalization (Figure 3). This trafficking defect can cause an alteration in transport efficiency of auxin leading to its intracellular accumulation at the apical cells, as shown for the *Δpinapinb* mutants where a reduced auxin export into the medium was reported (Viaene et al., 2014; Thelander et al., 2017), together with an accelerated differentiation of caulonemal cells, just as we observed for the *Δcnih2* mutant (Figure 2 A, C). Confirmation that the correct localization of PINA is dependent on its interaction with CNIH2 was obtained by generating C-terminal truncations of the CNIH2 protein (Figure 7A-B). This effect can be explained by the decreased protein-protein interaction that was observed between the truncated CNIH2-137 and PINA in the mbSUS assay (Figure 6B), and agree with previous reports demonstrating that the acidic domain in cornichon homologues from yeast and plants is important to establish a strong interaction with different cargo membrane proteins and necessary for their correct localization at the plasma membrane (Rosas-Santiago et al., 2017). Additionally, and may be particular for CNIH2, the presence of a potential phosphorylation residue (T149) seems to play an important role in the interaction with PINA, as evinced by the mislocalization of the transporter caused by removal of the T/S domain and the associated morphological changes caused in the protonema (Figure 7C), opening the possibility that the phosphorylated receptor could play a positive role in regulating cargo release to the plasma membrane.

We observed abnormal positioning of branching cells in the cornichon mutants (Figure 2A-B and Supplemental Figure S4). Until now, it has only been reported that mutants in Myosin VIII (Wu et al., 2011) and a Vapyrin-like (VPY-like) protein (Rathgeb et al., 2020) exhibit these branching defects. In moss protonemata, one of this cornichon-interacting proteins could be a VPY-like protein according to its localization to puncta in the cytoplasm and around the nucleus, coincident to that we observed for CNIH2 (Figure 5A). The localization of VPY and VPY-like proteins was denoted as vapyrin bodies related to the TGN and endosomes, but also associated with the endoplasmic reticulum (Pumplin et al., 2010; Feddermann et al., 2010; Liu et al., 2019; Bapaume et al., 2019; Rathgeb et al., 2020). Although the function of Vapyrin (VPY) and VPY-like proteins is unknown, they are only found in plants and possess a vesicle-associated protein (VAP) domain at their N-terminus, and several ankyrin repeat domains at the C-terminus, with both domains predicted to be involved in protein-protein interactions (Pumplin et al., 2010; Feddermann et al., 2010), opening the possibility that they could associate with CNIH2.

According to the interaction between CNIH1 and CNIH2 with PINA that we demonstrate in this work, it is possible that the phenotypes observed in the *Δcnih1, Δcnih2* and *Δcnih1Δcnih2* gametophores (Figure 2 E-F) might be due to an alteration in the intracellular concentration of auxins. Supporting this view are the bigger gametophores reported for the *Δpinb* mutant (Bennett et al., 2014), phenotype similar to that we observed for the *Δcnih2* mutant, confirming the role of CNIH2 as the cargo receptor for PINA and suggesting that PINB could also depend on a similar interaction for its incorporation into the plasma membrane.

The independence of PINA from CNIH1 to reach its apical location (Figure 3C) despite having observed the interaction of the two proteins (Figure 3 A-B), together with the pleiotropic effects generated by the single cornichon mutants (Figure 2), indicate that they may be caused by the mistargeting of additional membrane proteins as cornichon receptors interact and mediate the trafficking of many membrane proteins that pass through the secretory route (Herzig et al., 2012; Rosas-Santiago et al., 2015), including those related to cell proliferation and expansion, processes that are required for the formation of the gametophore shoot (Kawade et al., 2020).

The observation that CNIH2 is associated with Sec23G (Figure 5), one of the seven isoforms of Sec23 (Chang et al., 2021), leads us to propose that CNIH2 is located to a Sec23G structured ERES as part of the pathway that regulates PINA exit from the ER to the plasma membrane, and as an alternative trafficking mechanism in which the polarized distribution of PINA in moss cells is mainly mediated by the early secretory pathway. These results contrast with those reported for Arabidopsis where several processes like endocytosis, recycling and/or transcytosis, or the participation of the retromer complex and/or the exocyst complex have been proposed to establish the polarity to *At*PIN1 in Arabidopsis (Jaillais et al., 2007; Geldner et al., 2003; Drdová et al., 2013; Doyle et al., 2015). Our results expand our knowledge on the participation of different trafficking mechanisms of PIN proteins in different plant lineages.

## Materials and Methods

### Protein sequence analysis

Protein sequence alignments and phylogenetic analyses were obtained with the CLC Main Workbench v6 software (Qiagen). Phylogenetic tree was used from matrix pairwise data and using UPGMA algorithm, perform 100 bootstrap rounds.

For predicts Serine, Threonine and/or Tyrosine protein phosphorylation sites in PpCNIH1 and PpCNIH2 proteins, we employed the server NetPhos3.1 (http://www.cbs.dtu.dk/services/NetPhos/) (Blom et al., 1999).

### Moss strains, growth conditions and protoplast isolation

*Physcomitrium patens* (“Gransden” WT strain) was used in this study. Protonema was propagated routinely by spreading the tissue onto a cellophane disk laid inside a Petri dish with PpNH_4_ medium (1.03 mM MgSO_4_, 1.86 mM KH_2_PO_4_, 3.3 mM Ca(NO)_3_, 45 mM FeSO_4_, 2.72 mM (NH_4_)_2_-tartrate, 9.93 mM H_3_BO_4_, 220 nM CuSO_4_, 1.966 mM MnCl_2_, 231 nM CoCl_2_, 191 nM ZnSO_4_, 169 nM KI and 103 nM Na_2_MoO_4_) supplemented with 0.7% agar; the moss was grown at 24°C under 70 μE m^-2^ s^-1^ light and 16 h-light/ 8h-dark regime. The YFP-Golgi (YFP-*Gm*Man) line was proportionated by Dr. Magdalena Bezanilla (Dartmouth College, MA United States), *PpPINA_pro_:PpPINA-EGFP* line (refer as PINA-EGFP) was provided by Dra. Sundberg (Swedish University of Agricultural Science, Uppsala, Sweden).

For protoplasts isolation, seven-day old protonema tissue was digested in a 0.5% driselase and 8.5% Mannitol solution for one hour in a shaker at room temperature. Protoplasts were passed through one layer of Miracloth (Merck-Millipore) to remove undigested tissue and centrifuged 1,600 rpm for 7 min and washed in 8.5% Mannitol solution, three times. The protoplasts were resuspended in top agar (PpNH_4_ medium, 6% Mannitol, 10 mM CaCl_2_, 0.3% agar) or in liquid plating media (PpNH_4_ medium with 8.5% Mannitol, 10 mM CaCl_2_). Finally, for protonema regeneration, protoplasts were transferred into PRMB medium (PpNH_4_ medium supplemented with 6% Mannitol, 10 mM CaCl_2_ and 0.7% agar). We employed the protoplasts for moss transformation and as starting material for morphological evaluation.

### Plasmids

Almost all the entry and expression plasmids employed in this work were obtained from Dra. Magdalena Bezanilla and are compatible with the Gateway technology (Invitrogen, CA, USA). Plasmids for transient expression were: pDONR221 and pTZeoUBI-gate. Plasmids for generating gene mutation by CRISPR-Cas9 system, were: pENTR-PpU6P-L1L5r, L5L4, L4L3 and L3L2 and the pMZeo-Cas9-gate. For the generation of knock-in constructs by CRISPR-Cas9 and Homologous Directed Repair, the plasmids used were: pENTR-PpU6P-L1L2, pDONR-P1P4, pDONR-P3P2, pDONR-R4R3-3xmRuby-C, pDONR-R4R3-3xmNeon-C, pMZeo-Cas9-gate, pMH-Cas9-PpU6P-sgRNA-sec23g, pDONR-B1-Sec23G 5’ arm-B4, pDONR-B3-Sec23G 3’ arm-B2 and pGEM-gate. The plasmids employed to generate knock-out mutants by the Homologous Recombination technique were: pDONR-P1P4, pDONR-P3P2, pDONR-R4R3-loxP-Hygro-loxP and the pGEM-gate.

### Protospacers constructs for *CNIH1* CRISPR-Cas9 mutagenesis

To generate the *Cnih1* protospacers construct, the CRISPR-Cas9 system was employed, as described (Mallett et al., 2019). A protospacer was designed with the CRISPOR online software (crispor.tefor.net) using *P. patens* (Phytozome V11) as the genome and *S. pyogenes* (5’NGG3’) as the PAM parameters. Four protospacers were chosen based on high specificity scores and low off-target frequency along the *CNIH1* gene. Each protospacer was designed to target one of the four predicted exons of the *CNIH1* gene. Each protospacer and its reverse complement were synthesized as oligonucleotides adding the CCAT-sequence at the 5’ end of each to create sticky ends compatible with *BsaI* (Thermo Fisher, MA, USA). Each pair of oligonucleotides were annealed by PCR (500 pmol of each, 10 μl total volume, with the following setting conditions: 98°C 3 min, 0.1°C/s to oligo Tm, hold 10 min, 0.1°C/s to 25°C). In parallel, the pENTR-PpU6P-L1L5r, L5L4, L4L3 and L3L2 entry vectors were linearized with *BsaI* for 14-16 h at 37°C. Each protospacer was ligated with their respective entry vectors, using the Instant Sticky-end Ligation Master Mix (New England Biolabs, MA, USA), following manufacturer’s specifications, and generating pENTR-L1L5r-protospacer 1, pENTR-L5L4-protospacer 2, pENTR-L4L3-protospacer 3 and pENTR-L3L2-protospacer 4 constructs. The correct sequence of these entry constructs was confirmed by sequencing and finally recombined into the pMZeo-Cas9-gate expression vector by a four-fragment multisite Gateway recombination reaction (Invitrogen, CA, USA) with the LR clonase II plus, following manufacturer’s specifications. The pMZeo-Cas9/protospacers plasmid construct was transformed into moss protoplasts by the PEG transformation protocol.

### *CNIH2-3xmRuby* and *SEC23G-3xmNeon* tagging by CRISPR-Cas9 and Homologous Directed Repair (HDR) system

To generate the *CNIH2-3xmRuby* transgenic moss line, we employed the CRISPR-Cas9 system together with the Homologous Directed Repair system described in Mallet *et al*., 2019 (Mallett et al., 2019). To generate the pMZeo-Cas9/cn2 tag protospacer plasmid construct, first, a protospacer was designed and chosen with the parameters as described above. In this case the protospacer was chosen closest to the stop codon of the *CNIH2* gene. Then, the selected protospacer and its reverse complement were synthesized as oligonucleotides adding the CCAT-sequence at the 5’ end of each, to create sticky ends compatible with *BsaI* linearized entry vectors. The oligonucleotides were annealed by PCR and cloned into the pENTR-PpU6P-L1L2 entry vector as previously described; this construct was sent for sequencing and finally, it was recombined into the pMZeo-Cas9-gate expression vector by an LR clonase reaction following manufacturer’s specifications (Invitrogen, CA, USA).

To generate homology arms for tagging *CNIH2*, we amplified two fragments of 1,109 bp upstream of the start codon and 1,101 bp downstream of the *CNIH2* stop codon. The upstream and downstream fragments were cloned in the pDONR-P1P4 and pDONR-P3P2 vectors, respectively, using a BP clonase reaction. To generate the homology directed repair construct required for *CNIH2-3XmRuby* C-terminal gene tag, we used the three fragment Multisite Gateway cloning system (Invitrogen, CA, USA) and recombined the pDONR-B1 *-CNIH2* 5’ arm-B4, pDONR-R4R3-3xmRuby-C, pDONR-B3-C*NIH2* 3’ arm-B2 into the pGEM-gate destination vector by a triple LR reaction. The pMZeo-Cas9/*CNIH2* and pGEM-*CNIH2-3xmRuby* constructs were co-transformed into wild-type moss protoplasts.

To generate the *SEC23G-3xmNeon* transgenic moss line, the pMH-Cas9-PpU6P-sgRNA-*SEC23G* construct was used as protospacer plasmid. To produce the homology directed repair construct required for construction of the *SEC23G-3XmNeon* C-terminal gene tag, the pDONR-B1-*SEC23G*23 5’ arm-B4, pDONR-B3-*SEC23G* 3’ arm-B2 and pDONR-R4R3-3xmNeon-C constructs were used and recombined into the pGEM-gate destination vector by a triple LR reaction. pMH-Cas9-PpU6P-sgRNA-*SEC23G* and *pGEM-SEC23G-3xmNeon* constructs were finally co-transformed in protoplasts using the *CNIH2-3xmRuby* line as background.

### Generation of *PpCNIH2* knock-out constructs by Homologous Recombination

To obtain the *Δcnih2* null mutant, a replacement construct was generated, for this, a 1,196 bp PCR-amplicon upstream from the 5’ start site of the gene, and a 1,284 bp PCR-amplicon downstream from the stop codon, were amplified from genomic DNA and cloned independently by a BP clonase reaction into the pDONR-P1P4 and pDONR-P3P2 plasmids, respectively, according to manufacturer’s specifications (Invitrogen, CA, USA). The primers contained the appropriate *attb* sites. A *PmeI* restriction site was designed inserted at both 5’ and 3’ ends to linearize the knock-out construct. The entry clones were corroborated by sequencing. To generate the homologous DNA donor template flanking the hygromycin resistance cassette, the entry clones with homologous arms and the pDONR-R4R3-loxP-Hygro-loxP plasmid were recombined into the pGEM-gate plasmid in a three-fragment multisite Gateway recombination (Invitrogen, CA, USA), using an LR II clonase plus reaction. Finally, a *PmeI* digestion of the homologous DNA donor template was performed, the linearized construct was precipitated with ethanol before moss protoplast transformation.

For generation of the cornichon double mutant, the parental line *Δcnih1-23* was employed to create the *Δcnih2* null mutant by homologous recombination, in which the *CNIH2* gene was replaced by the hygromycin resistance cassette, as described above.

### Moss transformation and selection of transformants

For protoplast moss transformation, the Polyethylene Glycol-mediated (PEG) transformation protocol was used. For homologous recombination replacement, protoplasts were transformed with 15 or 30 μg of DNA. For the CRISPR-Cas9 mutagenesis, protoplasts were transformed with 15 μg of total DNA construct. For the CRISPR-Cas9 & HDR system protocol, protoplasts were co-transformed with 7.5 μg total of CRISPR-Cas9/protospacer plasmid construct and 7.5 μg of total homology plasmid. After transformation, plants were allowed to regenerate on PRMB for four days.

To select stable transformants of the *Δcnih2* null mutants, plants were moved to PpNH_4_ medium containing hygromycin (15 μg/ml). The potential knock-out transformants were cycled on and off in antibiotic plates for two 1-week intervals. For the selection of the CRISPR-Cas9 and the CRISPR-Cas9 & HDR transgenic lines, Zeocin (50 μg/ml) was employed. For the selection of the *SEC23G-3xmNeon* line, the transformants were grown on hygromycin containing medium. After plants were under selection for one week, the cellophane was changed to freshly PpNH_4_ medium three times every week. Finally, the surviving plants were picked with sterile tweezers and grown on PpNH_4_ medium (without cellophane) by 3-4 weeks to allow maximal growth for genomic DNA extraction.

### Genetic analyses

To identify and select the colonies transformed with the *CNIH2-3xmRuby* or *SEC23G-3XmNeon* constructs, internal primers 1463-Fw and 1462-Rv or s23G-Int-F and S23g-Int-R (Supplementary Table 1) were employed, respectively.

For the genotyping of potential *Δcnih1* single and double cornichon null mutants, primers were designed that allowed the amplification of 1,000 bp flanking the mutation target site of sgRNA’s to observe the differences in PCR band size in comparison with the wt DNA region. Potential mutant moss colonies were identified and screened using the primers 1268-Fw and 1512-Rv (Supplementary Table 1), and those in which the amplified PCR product was different from the WT size, were selected. The genomic DNA of two potential single and double mutant lines was isolated and confirmation of the PCR product was obtained by sequencing. *CNIH1* gene mutant expression analysis was corroborated by PCR amplification in the *Δcnih1* single and double mutant using the primers 252-Fw and 253-Rv (Supplementary Table 1).

For genotyping the potential *Δcnih2* null mutant line in single and double mutants, a PCR amplification was performed to synthesize the expected fragments of 5,331 bp for *CNIH2* genomic DNA and 6,035 bp for the knockout (hygromycin cassette) genomic DNA, using the primers 327-Fw and 328-Rv (Supplementary Table 1). Confirmation of the replacement of the *CNIH2* gene by the hygromycin cassette was obtained by PCR analyses. In one reaction, using the primers 327-Fw and 319-Rv (Supplementary Table 1), a 3,059 bp PCR fragment corresponding to the 5’ homologous genomic region and the hygromycin cassette sequence was obtained, as expected. By using the primers 320-Fw and 328-Rv (Supplementary Table 1), a second 4,120 bp PCR fragment corresponding to the 3’ homologous genomic region and the hygromycin cassette sequence was also obtained, as anticipated. Absence of the *CNIH2* transcript in the *Δcnih2* single and double mutants was confirmed by PCR amplification using the primers 298-Fw and 299-Rv (Supplementary Table 1).

### DNA extraction by the CTAB method

For genomic DNA extraction, the CTAB (cetyl trimethylammonium bromide) method was employed, with the following modifications. A DNA extraction Buffer (0.1 M Tris-HCl pH 8.0, 1.4 M NaCl, 2.0% CTAB, 20 mM Na2EDTA, 2.0% PVP-40) was previously prepared adding β-mercaptoethanol (10 mM) and ascorbic acid (0.1%). After that, half of a moss colony was taken with sterile tweezers, and to remove excess water, the tissue was squeezed between several sheets of sterile filter paper (Whatman 3M). The dried tissue was collected in a 1.5 ml sterile tube and immediately frozen in liquid N2. The tissue was homogenized in a prechilled microcentrifuge pestle, adding 200 μl of DNA extraction Buffer previously pre-warmed to 65°C and ground until a green liquid mix was formed. Then, 1 μl RNAse A (10 mg/ml) was added, and the tubes were incubated at 65°C for 5 min. After that, 100 μl of the mix chloroform:isoamyl alcohol (24:1) was added to the tubes and mixed thoroughly; the phases were separated by centrifugation at 12,000 rpm for 10 min. The upper phase was collected to precipitate the DNA by adding an equal volume of Isopropanol and mixed by vortex; followed by incubation at −20°C for at least, 15 min. Working in a sterile hood, the supernatant was discarded, and the pellet was washed with 70% Ethanol and air dried. Finally, the pellet was dissolved in 25 μl sterile ddH2O.

### RNA extraction and cDNA synthesis

For extraction of total RNA from protonema tissue to genotype the cornichon single and double mutants, Plant RNA purification Reagent (Invitrogen, CA, USA) was used, followed by DNAse I treatment (Thermo Fisher Scientific, MA, USA) according to manufacturer’s recommendations. For the analysis of cornichon gene expression, synthesis of cDNA was performed using an oligo(dT) primer and RevertAid M-MuLV reverse transcriptase (Thermo Fisher Scientific, MA, USA) following manufacturer’s protocol.

### Morphological analysis

For protonema morphological analysis, protoplasts obtained from 7-day protonemal tissue were resuspended in 0.5 μl liquid plating medium (PpNH_4_ medium supplemented with 8.5% Mannitol and 10 mM CaCl_2_) and plated in PRMB medium plates (PpNH_4_ medium supplemented with 6% Mannitol, 10 mM CaCl_2_ and 0.7 % Agar) with cellophane overlays for four days, and then, transferred to PpNH_4_ medium for four days. After this, the protonema tissue was mounted onto a microscope slide with PpNH_4_ liquid medium and covered with a cover slide. Tissue was observed under an inverted microscope (Nikon, Eclipse) and images were acquired with a digital camera (Nikon 7500). Calcofluor was used to enhanced cell morphology by staining the cell wall and observed in a fluorescence stereomicroscope (Zeiss Axioscope) with an objective of 10X; images were obtained with a CCD camera (Photometrics CoolSnapcf Monochromatic) and analyzed with the software ImageJ/Fiji (Schindelin et al., 2012). Images from every line were adjusted to sharpen and contrast to improve the division cell lines. Quantification of caulonema/cloronema ratio was made manually.

Gametophore morphological analysis was performed in a colony assay. Seven-day-old protonema tissue sections of 3×3 mm were placed carefully with forceps on PpNH_4_ solid medium and were grown under 24°C 16h-light/8h-dark photoperiod for four weeks. The gametophores with more than five fillids and well-formed rhizoids (under stereomicroscope observation) were collected and counted. The moss colony and gametophores images were taken with a digital camera (Olympus or Sony α).

### Protonema sample preparation for Brefeldin-A treatment

For Brefeldin A (BFA; Sigma-Aldrich, MO, USA) treatment, several pieces of cellophane containing a one-week-old protonema were carefully cut with a scalpel, placing them on PpNH_4_ medium plates supplemented with 50 μM of BFA or DMSO (control plates). Plates were left in incubation for 24 h at 24°C (16h/8h light/dark regime). For confocal microscopy observation of protonema tissue, a piece of cellophane containing the protonema was cut and placed face down onto an agar pad containing 50 μl of Hoagland’s medium (4mM KNO_3_, 2mM KH_2_PO_4_, 1mM Ca(NO_3_)_2_, 89μM Fe citrate, 300μM MgSO_4_, 9.93μM H_3_BO_3_, 220nM CuSO_4_, 1.966μM MnCl_2_, 231nM CoCl_2_, 191nM ZnSO_4_, 169nM KI, 103nM Na_2_MoO_4_) with 1% agar and 1% sucrose. The cellophane was removed by sliding it carefully, leaving the tissue attached to the agar pad. Ten μl of liquid Hoagland’s medium and 1% sucrose were added onto the tissue, covered with a coverslip, and sealed with 1:1:1 Vaseline:Lanolin:Paraffin mixture. To maintain the tissue under exposure to BFA during observation, 50 μM BFA was added to the mix of liquid Hoagland’s medium.

### Confocal Microscopy and Co-localization assay

For *CNIH2-3xmRuby* fluorescence was observed in a confocal microscope LSC Nikon Ti2, employing an excitation wavelength of 561 and an emission of 647nm with a 1.49 NA, Apo 60x immersion oil objective. The fluorescence of the YFP-Golgi marker was obtained by exciting at 488 nm (0.5% laser power) and observed at 524 nm, with a 60X oil immersion objective with a N.A. 1.3 using an inverted confocal microscope Olympus FV1000. For both transgenic lines, optical sections of 0.75 to 1.00 μm in -z were taken for acquisition of images. The Images were viewed with ImageJ (Schindelin et al., 2012).

For co-localization analysis between SEC23G and CNIH2 an inverted confocal microscope Olympus FV1000 was employed. Fluorescence from *SEC23G-3XmNeon* was obtained by exciting at 488 nm (0.5% power laser) and observed at an emission of 510 nm. *CNIH2-3xmRuby* fluorescence was exciting at 543 nm (5% power laser) and observed at 655 nm emission in a 60X oil immersion objective with a N.A 1.3. Confocal images were taken every 0.75 to 1.00 μm in z axis. The Images were viewed and analyzed in ImageJ. For co-localization analysis, the Co-loc2 plugin in Image-J was employed to derive a Pearson’s correlation coefficient; values above 0.5 are indicative of colocalization.

Dot size quantification was made in ImageJ by sharpening and thresholding three different optical confocal images from *CNIH-3xmRuby*, *SEC23G-3xmNeon* and YFP-Golgi lines, respectively. Vesicles were selected with wand tool, the area of a total of 35 random vesicles was measured and plotted.

### Mating-based split ubiquitin yeast system

For the detection of protein-protein interactions, we employed the *m*ating-*b*ased *S*plit *U*biquitin *S*ystem (mbSUS) in yeast cells (Obrdlik et al., 2004; Lalonde et al., 2010). Plasmid constructs were generated using the coding sequence for *PINA*, *CNIH1*, CNIH*2, CNIH2-141* and *CNIH2-137* without the stop codon and cloned into the pDONR221 plasmid by a BP reaction (Invitrogen, CA, USA). Once we verified by sequencing the correct cloned genes, we used an LR clonase reaction (Invitrogen, CA, USA) to transfer the *PINA* gene to the pMETYC_GW (Cub clones) and *CNIH1, CNIH2, CNIH2-141* and *CNIH2-137* to the pXN32_GW (Nub clones) vectors, respectively. For the mbSUS assay, yeast media were prepared as previously described in (Lalonde et al., 2010; Jones et al., 2014). The THY.AP4 (*MATa ura3, leu2, lexA::LacZ::trp1 lexA::HIS3 lexA::ADE2*) and THY.AP5 (*MATa URA3, leu2, trp1, his3 loxP::ade2*) yeast strains were transformed with the pMETYC_GW and pNX32_GW vectors, respectively. Yeast cells were transformed by the LiAc treatment.

### Bi-Fluorescence Complementation assay in leaf epidermal cells from *Nicotiana benthamiana*

Complementation of EYFP fluorescence (BiFC) experiments were done according to Rosas-Santiago et. al, (2015)(Rosas-Santiago et al., 2015). To obtain the expression clone for each gene employed in the BiFC assays, an LR gateway-based recombination reaction (Invitrogen, CA, USA) with either pYFC43 or pYFN43 was carried out. Leaves were infiltrated with the constructs *pYFC43-PpPINA,pYFC43-AtPIP2A,pYFN43-PpCNIH1*,*pYFN43-PpCNIH2* and *pYFN43-AtPIP2A. Agrobacterium tumefaciens* GV3101 strain cells were transformed with each construct by electroporation and grown in 30 ml of LB medium with rifampicin (50 μg ml^−1^) and spectinomycin (50 μg ml^−1^) or kanamycin (50 μg ml^−1^) at 28 °C at an OD_600_ of 0.3 to 0.5. Leaves were infiltrated with a bacterial culture with an OD_600_ of 0.3 resuspended in sodium phosphate buffer pH 7.0, 0.1 mM acetosyringone (Sigma-Aldrich, MO, USA) and 28 mM glucose.

### Changes in subcellular localization of PINA-EGFP in moss protonema and complementation assays

To observe changes in PINA subcellular location in cornichon single mutants, we employed the reporter line *PpPINApro:PpPINA-EFGP*. *Δcnih1* single mutant was obtained by the CRISPR-Cas9 technique, while the *Δcnih2* single mutant was obtained by homologous recombination, as previously described. Selected single cornichon mutants were picked and observed under a confocal microscope to identify the location of the PINA-EFGP fluorescence in protonema apical cells. A piece of cellophane containing 7-day old protonema was cut and placed onto a microscope slide containing 20 μl of PpNH_4_ liquid medium and covered with a coverslip and sealed with nail polish for observation of EGFP (excitation and emission wavelengths of 488 nm and 510 nm, respectively with a 0.5% of laser power in an inverted confocal microscopy (Olympus FV1000).

To clone the *CNIH2* wildtype gene and the C-terminus truncated versions *CNIH2-141* and *CNIH2-137*, total RNA was obtained from *P. patens* protonemal tissue using the Plant RNA purification Reagent (Invitrogen, CA, USA), followed by a DNAse I treatment (Thermo Fisher Scientific, MA, USA) following manufacturer’s recommendations. Synthesis of cDNA was performed using an oligo(dT) primer and RevertAid M-MuLV reverse transcriptase (Thermo Fisher Scientific, MA, USA) following manufacturer’s protocol. The predicted coding sequences for *CNIH2, CNIH2-141* and *CNIH2-137* were amplified by PCR with the appropriate *attb sites* (B1 and B2) and each PCR product was cloned in the pDONR-221 entry vector by a BP clonase reaction. These constructs then were cloned in pMZeo-Ubi-gate expression plasmid by an LR reaction. The entry and the expression clones were verified by restriction enzyme analysis and by sequencing at the sequencing unit of Instituto de Biotecnología, UNAM, México.

For complementation assays, each construct was transformed into protoplasts for transient expression in the moss single mutant *Δcnih2-3/ PpPINA-EFGP*. After transformation, plants were allowed to regenerate on PRMB medium for four days and then were maintained in zeocin selection for 12 days before were observed under a confocal microscope as describe above in this section. Each image corresponding to each complementation with CNIH2 and C-terminus truncated versions were z-projection and changed to 8-bit image for measure the intensity of PINA-EGFP fluorescence (0-255), after was delimited an area of 46.8 μm2 with a polygon ROI at apex zone of protonema cell.

### Statistical analysis

The data are presented as bars and box-plots obtained with Origin software (OriginLab). All statistical analysis were performed using Excel. Student’s *t* test for unpaired data with equal variance was used. For gametophore analysis ANOVA and Tukey-Kramer post hoc test was performed. P values greater than 0.05 were reported as not significant of ns. P values equal or smaller than 0.05 or 0.001 were reported as significant and highly significant, respectively.

### Accession numbers

Plant and algae cornichon homologue sequences were obtained from ARAMEMNON (http://aramemnon.botanik.uni-koeln.de/), Phytozome (https://phytozome-next.jgi.doe.gov/) and EnsemblPlants for algae Chara braunii (https://plants.ensembl.org/Chara_braunii/Info/Index). Names and accession number are as follows: *Arabidopsis thaliana* (AtCNIH1: At3g12180.1, AtCNIH2: At1g12340.1 AtCNIH3: At1g62880.1, AtCNIH4: At1g12390.1, AtCNIH5: At4g12090.1), *Oryza sativa* (OsCNIH1: Os06g04500.1 OsCNIH2: Os12g32180.1*, Zea mays* (ZmCNIH1: GRMZM2G073023.01, ZmCNIH2: GRMZM2G018885.01 and ZmCNIH3: GRMZM2G124658.01), *Populus trichocarpa* (PtCNIH4: Potri001g116100.1, PtCNIH3: Potri003g116400.1, PtCNIH1: Potri006g057300.3, and PtCNIH2: Potri016g051000.1), *Ananas comosus* (AcCNIH: Aco015328.1), *Zostera marina* (ZmaCNIH1: Zosma28g00840, ZmaCNIH3: Zosma*42g*01060 and ZmaCNIH2: Zosma153g00310), *Selaginella moellendorfi* (SmCNIH: 93931(PAC:15402723)), *Physcomitrium patens* (PpCNIH1: *Pp3c11_17020V3.3* and PpCNIH2: *Pp*3c7_11500V3.3), *Marchantia polymorpha* (MpCNIH: *Mapoly0124s0019), Chlamydomonas reinhardtii* (CrCNIH: Cre01.g036550_4532), *Dunaliella salina* (DsCNIH: Dusal.0011s00016), and *Chara braunii* (CbCNIH: GBG61058.1), Fungi cornichon homologue sequences were obtained from Yeast Genome database (www.yeastgenome.org) and NCBI: *Saccharomyces cerevisiae* (ScErv14p: SGD:S000003022), *Schizosaccharomyces pombe* (SpCNIH: NP_594657.1), *Neurospora crassa* (NcCNIH: XP_011395262.1) and *Aspergillus nidulans* (AnCNIH: XP_662799.1).

## Acknowledgments

Carolina Yáñez-Domínguez is a doctoral student from the Programa de Doctorado en Ciencias Biomédicas, Universidad Nacional Autónoma de México (UNAM) and received a CONACyT fellowship número de becario 662829. This report was supported by Grant 2041 from CONACYT, Mexico to O.P. We would like to thank past and actual members of Bezanilla’s lab: Dr. Mingqin Chang, Dr. Carlisle Bascom Jr., Jackie O’Sullivan and Samantha Ryken for technical support. We also thank Dr. Xiaohang Cheng for helpful CRISPR-Cas9 and CRISPR-Cas&HDR assistance and Dr. Shuzon Wu for help with BFA treatment and imaging. We thank Dr. Eva Sundberg and Dr. Katarina Landberg for kindly facilitating us the PpPINA-EGFP moss line. We thank Guadalupe Muñoz García for technical support. We acknowledge Laboratorio Nacional de Microscopía Avanzada (LNMA, IBt UNAM).

## Author contributions

Conceptualization: CYD, OP

Methodology: CYD, DLG, DMTC

Investigation: CYD, DLG, DMTC

Funding acquisition: OP

Writing – original draft: CYD, OP

Writing – review & editing: CYD, DLG, DMTC, MB, OP

