## Supplemental Information for "A cornichon protein controls polar localization of the PINA auxin transporter in *Physcomitrium patens*"

Figure S1

A

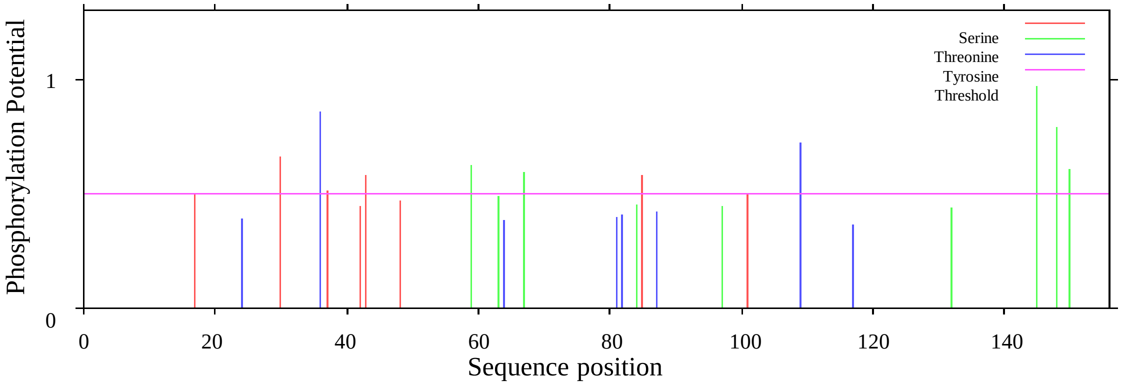


B

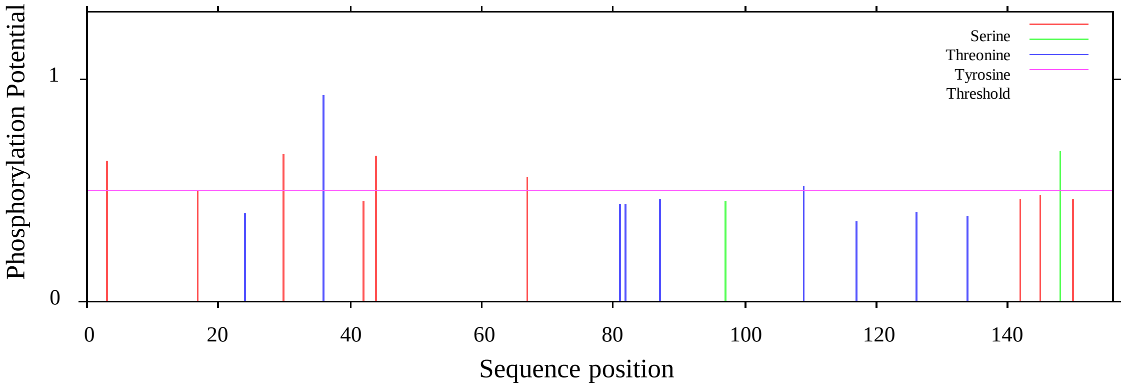


C

| | | 1 | 2 | 3 | 4 | 5 | 6 | 7 | 8 | 9 | 10 | 11 | 12 | 13 | 14 | 15 | 16 | 17 | 18 | 19 | 20 | 21 | 22 | 23 | 24 | 25 | 26 | 27 | 28 | 29 |
|---|---|---|---|---|---|---|---|---|---|---|---|---|---|---|---|---|---|---|---|---|---|---|---|---|---|---|---|---|---|---|
| AtCNIH4 | 1 | | 81.75 | 70.29 | 69.57 | 56.93 | 58.39 | 57.25 | 53.24 | 46.79 | 42.95 | 46.67 | 51.08 | 47.68 | 45.03 | 46.21 | 45.03 | 44.81 | 40.56 | 53.85 | 40.41 | 36.49 | 33.33 | 33.33 | 30.71 | 30.99 | 30.99 | 24.82 | 27.22 | 24.53 |
| AtCNIH3 | 2 | 112 | | 63.04 | 65.94 | 54.01 | 54.74 | 55.07 | 51.80 | 46.15 | 41.67 | 46.67 | 48.92 | 46.36 | 45.03 | 43.45 | 45.70 | 40.26 | 42.66 | 51.75 | 38.36 | 37.16 | 35.37 | 34.75 | 34.29 | 32.39 | 32.39 | 25.53 | 27.85 | 26.42 |
| PtCNIH4 | 3 | 97 | 87 | | 93.48 | 62.32 | 62.32 | 58.99 | 55.71 | 42.95 | 41.03 | 45.03 | 48.57 | 43.42 | 42.11 | 43.15 | 45.39 | 41.29 | 39.58 | 46.53 | 41.10 | 38.51 | 35.37 | 33.10 | 30.50 | 33.57 | 35.66 | 25.35 | 24.68 | 25.79 |
| PtCNIH3 | 4 | 96 | 91 | 129 | | 63.04 | 63.04 | 60.43 | 52.14 | 44.23 | 42.31 | 46.36 | 47.14 | 44.74 | 42.76 | 42.47 | 46.71 | 40.65 | 38.19 | 47.92 | 42.47 | 40.54 | 36.73 | 31.69 | 29.08 | 32.17 | 32.87 | 25.35 | 24.68 | 25.79 |
| OsCNIH1 | 5 | 78 | 74 | 86 | 87 | | 93.33 | 63.77 | 58.99 | 44.23 | 44.87 | 49.33 | 47.10 | 44.37 | 43.71 | 45.52 | 43.05 | 41.56 | 41.26 | 43.66 | 40.41 | 38.78 | 36.30 | 32.86 | 31.43 | 29.58 | 30.99 | 31.21 | 26.92 | 26.42 |
| ZmCNIH1 | 6 | 80 | 75 | 86 | 87 | 126 | | 63.77 | 56.12 | 42.95 | 43.59 | 48.67 | 46.38 | 42.38 | 42.38 | 44.83 | 41.72 | 38.96 | 39.86 | 45.77 | 39.73 | 37.41 | 34.25 | 32.14 | 30.71 | 30.28 | 30.99 | 33.33 | 28.21 | 28.30 |
| ZmaCNIH1 | 7 | 79 | 76 | 82 | 84 | 88 | 88 | | 63.50 | 42.95 | 39.74 | 46.62 | 48.53 | 46.67 | 45.33 | 45.45 | 47.33 | 41.83 | 38.03 | 44.29 | 42.18 | 36.49 | 35.37 | 32.86 | 29.50 | 31.91 | 31.91 | 29.50 | 24.53 | 27.50 |
| ZmaCNIH3 | 8 | 74 | 72 | 78 | 73 | 82 | 78 | 87 | | 43.31 | 38.85 | 42.95 | 48.89 | 45.03 | 42.38 | 44.37 | 44.37 | 40.26 | 41.96 | 42.14 | 39.46 | 35.81 | 33.78 | 34.51 | 32.86 | 34.51 | 35.92 | 33.57 | 23.12 | 26.88 |
| PpCNIH2 | 9 | 73 | 72 | 67 | 69 | 69 | 67 | 67 | 68 | | 71.79 | 61.15 | 52.87 | 48.41 | 49.04 | 47.13 | 49.04 | 48.41 | 51.59 | 39.74 | 39.49 | 40.13 | 38.85 | 32.69 | 33.12 | 33.76 | 34.39 | 31.85 | 26.42 | 26.88 |
| PpCNIH1 | 10 | 67 | 65 | 64 | 66 | 70 | 68 | 62 | 61 | 112 | | 56.05 | 45.86 | 43.95 | 43.95 | 42.68 | 47.13 | 43.95 | 44.59 | 32.69 | 38.22 | 39.49 | 37.58 | 29.49 | 29.94 | 31.21 | 30.57 | 26.75 | 25.79 | 24.38 |
| MpoCNIH | 11 | 70 | 70 | 68 | 70 | 74 | 73 | 69 | 64 | 96 | 88 | | 57.43 | 52.67 | 52.67 | 50.00 | 56.67 | 50.98 | 49.66 | 43.24 | 45.75 | 43.51 | 41.83 | 35.81 | 37.84 | 35.81 | 35.14 | 29.73 | 25.00 | 28.75 |
| SleCNIH | 12 | 71 | 68 | 68 | 66 | 65 | 64 | 66 | 66 | 83 | 72 | 85 | | 48.00 | 45.33 | 49.65 | 50.00 | 43.14 | 50.70 | 49.26 | 40.82 | 39.46 | 37.67 | 35.46 | 37.68 | 35.25 | 37.41 | 30.66 | 29.56 | 26.25 |
| OsCNIH2 | 13 | 72 | 70 | 66 | 68 | 67 | 64 | 70 | 68 | 76 | 69 | 79 | 72 | | 87.25 | 70.47 | 71.81 | 60.53 | 45.70 | 42.00 | 45.75 | 38.31 | 40.13 | 30.87 | 31.54 | 30.20 | 33.56 | 28.67 | 29.56 | 29.56 |
| ZmCNIH2 | 14 | 68 | 68 | 64 | 65 | 66 | 64 | 68 | 64 | 77 | 69 | 79 | 68 | 130 | | 74.50 | 69.13 | 61.84 | 46.36 | 40.00 | 47.06 | 38.31 | 39.47 | 30.87 | 32.89 | 29.53 | 32.21 | 28.67 | 28.30 | 30.19 |
| ZmCNIH3 | 15 | 67 | 63 | 63 | 62 | 66 | 65 | 65 | 63 | 74 | 67 | 74 | 70 | 105 | 111 | | 59.06 | 53.95 | 42.07 | 36.11 | 43.62 | 36.42 | 35.57 | 31.47 | 29.37 | 31.47 | 34.97 | 28.47 | 26.42 | 27.04 |
| AcoCNIH | 16 | 68 | 69 | 69 | 71 | 65 | 63 | 71 | 67 | 77 | 74 | 85 | 75 | 107 | 103 | 88 | | 63.82 | 50.99 | 40.67 | 50.33 | 41.56 | 40.79 | 33.56 | 36.24 | 30.87 | 33.56 | 30.67 | 31.45 | 30.19 |
| ZmaCNIH2 | 17 | 69 | 62 | 64 | 63 | 64 | 60 | 64 | 62 | 76 | 69 | 78 | 66 | 92 | 94 | 82 | 97 | | 43.51 | 39.87 | 44.87 | 36.94 | 35.48 | 26.97 | 28.29 | 26.32 | 29.61 | 24.84 | 27.04 | 27.04 |
| CbCNIH | 18 | 58 | 61 | 57 | 55 | 59 | 57 | 54 | 60 | 81 | 70 | 74 | 72 | 69 | 70 | 61 | 77 | 67 | | 34.48 | 38.10 | 36.49 | 34.69 | 33.80 | 35.21 | 34.51 | 36.62 | 34.51 | 30.62 | 28.75 |
| AtCNIH5 | 19 | 77 | 74 | 67 | 69 | 62 | 65 | 62 | 59 | 62 | 51 | 64 | 67 | 63 | 60 | 52 | 61 | 61 | 50 | | 37.58 | 33.56 | 31.76 | 27.27 | 26.24 | 26.06 | 25.35 | 22.14 | 24.68 | 26.25 |
| AtCNIH1 | 20 | 59 | 56 | 60 | 62 | 59 | 58 | 62 | 58 | 62 | 60 | 70 | 60 | 70 | 72 | 65 | 77 | 70 | 56 | 56 | | 47.97 | 47.62 | 31.51 | 30.82 | 24.66 | 26.03 | 27.89 | 21.38 | 25.16 |
| PtCNIH1 | 21 | 54 | 55 | 57 | 60 | 57 | 55 | 54 | 53 | 63 | 62 | 67 | 58 | 59 | 59 | 55 | 64 | 58 | 54 | 50 | 71 | | 87.76 | 28.38 | 29.05 | 27.70 | 25.68 | 28.38 | 27.04 | 25.62 |
| PtCNIH2 | 22 | 49 | 52 | 52 | 54 | 53 | 50 | 52 | 50 | 61 | 59 | 64 | 55 | 61 | 60 | 53 | 62 | 55 | 51 | 47 | 70 | 129 | | 26.71 | 27.40 | 27.40 | 26.03 | 26.53 | 25.95 | 25.16 |
| ScErv14 | 23 | 47 | 49 | 47 | 45 | 46 | 45 | 46 | 49 | 51 | 46 | 53 | 50 | 46 | 46 | 45 | 50 | 41 | 48 | 39 | 46 | 42 | 39 | | 72.14 | 61.43 | 59.29 | 43.97 | 24.84 | 23.90 |
| CaCNHI | 24 | 43 | 48 | 43 | 41 | 44 | 43 | 41 | 46 | 52 | 47 | 56 | 52 | 47 | 49 | 42 | 54 | 43 | 50 | 37 | 45 | 43 | 40 | 101 | | 68.35 | 67.63 | 46.38 | 24.53 | 26.42 |
| AnCNHI | 25 | 44 | 46 | 48 | 46 | 42 | 43 | 45 | 49 | 53 | 49 | 53 | 49 | 45 | 44 | 45 | 46 | 40 | 49 | 37 | 36 | 41 | 40 | 86 | 95 | | 86.96 | 46.76 | 25.79 | 22.64 |
| NcCNHI | 26 | 44 | 46 | 51 | 47 | 44 | 44 | 45 | 51 | 54 | 48 | 52 | 52 | 50 | 48 | 50 | 50 | 45 | 52 | 36 | 38 | 38 | 38 | 83 | 94 | 120 | | 46.04 | 24.53 | 22.01 |
| SpCNHI | 27 | 35 | 36 | 36 | 36 | 44 | 47 | 41 | 47 | 50 | 42 | 44 | 42 | 43 | 43 | 41 | 46 | 38 | 49 | 31 | 41 | 42 | 39 | 62 | 64 | 65 | 64 | | 27.50 | 25.62 |
| CreCNHI | 28 | 43 | 44 | 39 | 39 | 42 | 44 | 39 | 37 | 42 | 41 | 40 | 47 | 47 | 45 | 42 | 50 | 43 | 49 | 39 | 34 | 43 | 41 | 39 | 39 | 41 | 39 | 44 | | 37.74 |
| DsCNHI | 29 | 39 | 42 | 41 | 41 | 42 | 45 | 44 | 43 | 43 | 39 | 46 | 42 | 47 | 48 | 43 | 48 | 43 | 46 | 42 | 40 | 41 | 40 | 38 | 42 | 36 | 35 | 41 | 60 | |

**Figure S1. *In silico* analysis of putative phosphorylation sites of moss cornichon proteins and proteins pairwise comparison matrix. A)** Predicted phosphorylation sites for CNIH1 protein. **B)** Predicted phosphorylation sites for CNIH2 protein. Color code: Serine in red, Threonine in green and Tyrosine in blue, NetPhos3.1 server prediction was used for the in silico analysis. **C)** Pairwise comparison matrix for the cornichon proteins from plants, algae, and fungi representative organisms. Upper right corner values correspond to protein identity percentage, while the lower-left corner shows the number of identical amino acids. The data were analyzed with the software CLC Main Workbench 8.1.

Figure S2

**A**

sgRNA's design to knock-out cnih1 gene:

a) CN1-1E-sgRNA 5' TTCGCTGTTGTCTCGCTGCT 3'
**b) CN1-2E-sgRNA 5' TAAGAAGGTAGGTGCAGCCC 3'**
**c) CN1-3E-sgRNA 5' AGGTGACTGAAGATCTCCGT 3'**
d) CN1-4E-sgRNA 5' CCTCATGCTCCAGAACTAGG 3'

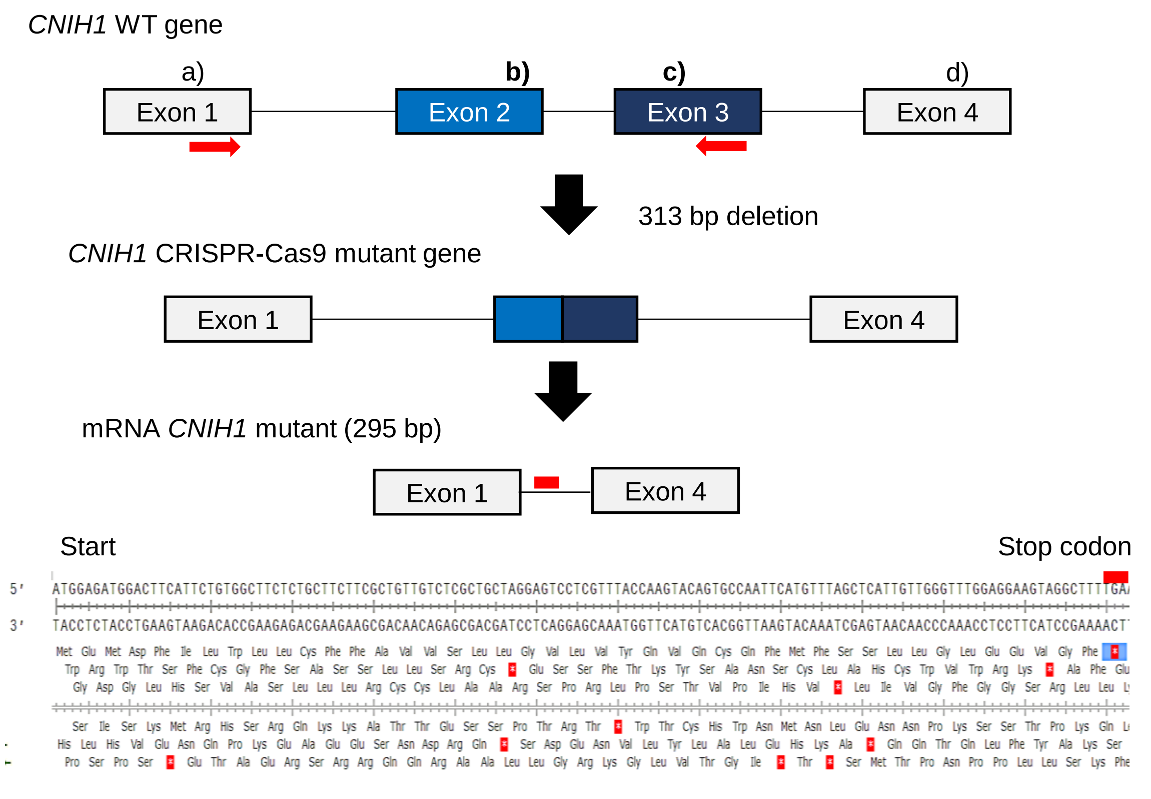


**B**

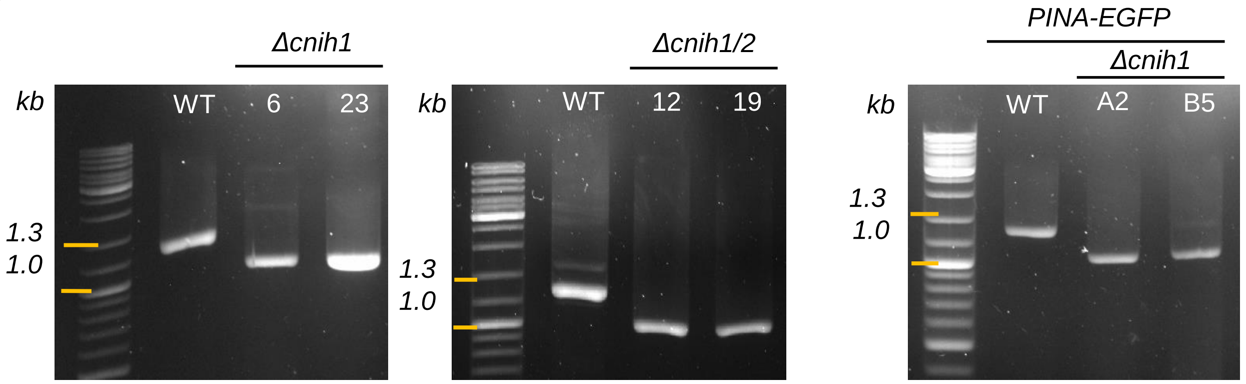


**C**

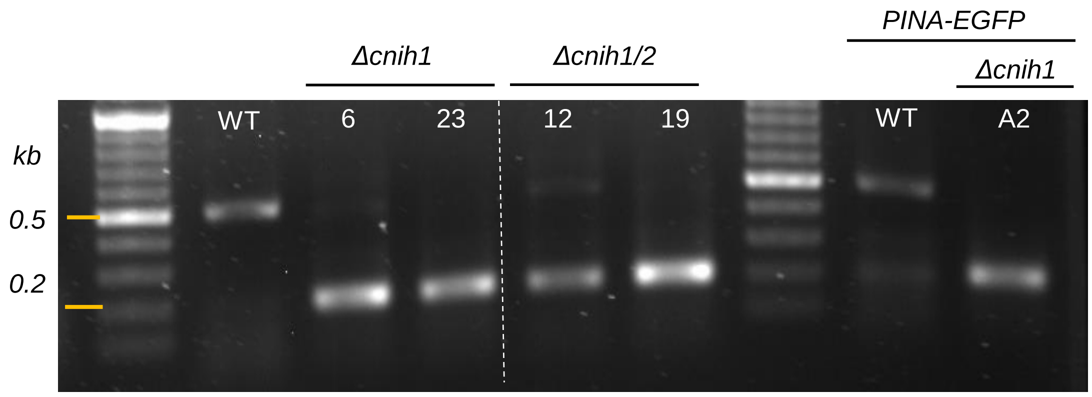

**Figure S2. Generation of *Δcnih1* mutant lines by the CRISPR-Cas9 system. A)** Schematic strategy for mutation of the *CNIH1* gene by the CRISPR-Cas9 system. Four different sgRNA were made (a to d); each sgRNA targeted one of the four exons of the *CNIH1* gene. Only the b and c sgRNA's (bold type) were efficient and deleted a total of 313 bp (removing the second intron and part of Exon 2 and Exon 3) resulting in the *Δcnih1* mutant line. This mutation generated an in-frame premature stop codon at nucleotide 132 (indicated by a red line) that codifies for 43 amino acids (predicted peptide size of 5 kDa). **B)** Comparison of the PCR products from the WT *CNIH1* gene (1,309 bp); the single *Δcnih1* mutant lines (1,000 bp) (#6 and #23) and double *Δcnih1/2* mutant lines (#12, #19) in WT parental line, and in the reporter PINA-EGFP lines (A2, B5). Amplified PCR bands from genomic DNA extractions. **C)** Comparison between WT *CNIH1* and *Δcnih1* single mutant coding sequences (cDNA). Agarose DNA gel (1%) shows amplified PCR bands from cDNA of WT *CNIH1* (468 bp); *Δcnih1* single mutant lines (295 bp) (#6 and #23) and double *Δcnih1/2* mutant lines (#12 and #19), and in the PINA-EGFP line *Δcnih1* single mutant (A2).

Figure S3

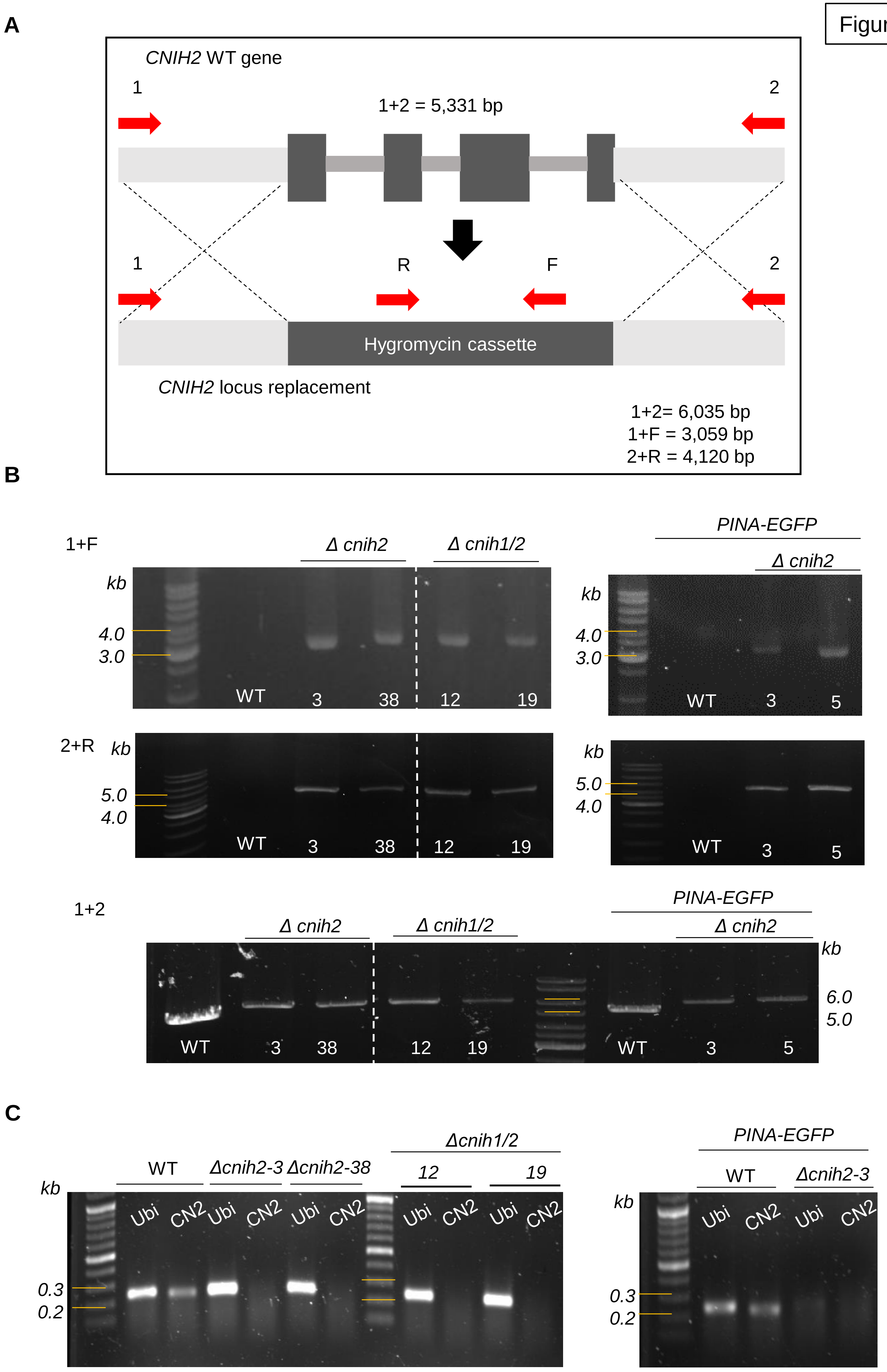
A
CNIH2 WT gene
1
2
1+2 = 5,331 bp
1
R
F
2
Hygromycin cassette
CNIH2 locus replacement
1+2= 6,035 bp
1+F = 3,059 bp
2+R = 4,120 bp
B
1+F
Δ cnih2
Δ cnih1/2
kb
4.0
3.0
WT
3
38
12
19
PINA-EGFP
Δ cnih2
kb
4.0
3.0
WT
3
5
2+R
kb
5.0
4.0
WT
3
38
12
19
kb
5.0
4.0
WT
3
5
1+2
Δ cnih2
Δ cnih1/2
PINA-EGFP
Δ cnih2
kb
6.0
5.0
WT
3
38
12
19
WT
3
5
C
WT
Δcnih2-3
Δcnih2-38
Δcnih1/2
12
19
kb
Ubi
CN2
Ubi
CN2
Ubi
CN2
Ubi
CN2
Ubi
CN2
0.3
0.2
PINA-EGFP
WT
Δcnih2-3
kb
Ubi
CN2
Ubi
CN2
0.3
0.2

**Figure S3. Generation of *Δcnih2* mutant lines and their genotypification. A)** *CNIH2* disruption strategy showing the genomic location of *CNIH2* locus and primers (red arrows) for genetic analyses. **B)** Agarose DNA gel (1%) showing the presence and the replacement by the hygromycin cassette by PCR amplified products from WT and *Δcnih2* single mutant lines (#3 and #38), *Δcnih1/2* double mutants (#12 and #19), and *Δcnih2* single mutant lines (#3 and #5) in the PINA-EGFP genetic line. **C)** Agarose DNA gel (1%) shows amplified PCR bands from cDNA of *CNIH2* (468 bp) in WT and in PINA-GFP lines, in comparison with the absence of *CNIH2* mRNA transcript in the *Δcnih2* single mutant lines (#3 and #38), *Δcnih1/2* double mutant lines (#12 and #19) and *Δcnih2* /PINA-GFP line (#3).

Figure S4

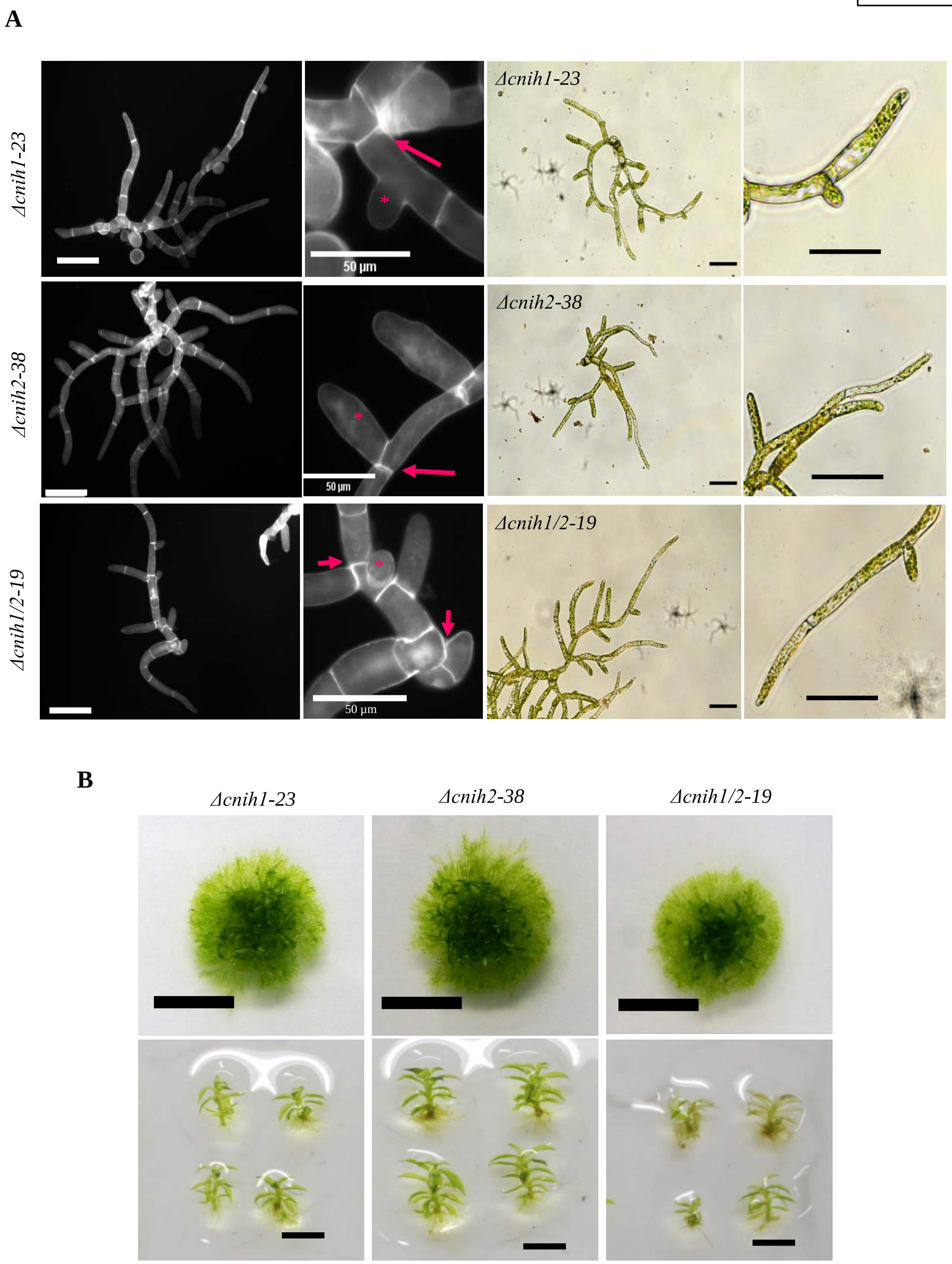


**Figure S4. Cornichon mutants have pleiotropic effects during the life cycle. A)** (Left panel) Protonema from WT and cornichon mutants stained with Calcofluor White after 7 d growth, visualized in an epifluorescence microscope; scale = 100 µm. Insets shows cell division (arrow) and lateral initial branch cell (*). (Right panel) Brightfield images of seven-day-old protonema from WT and cornichon mutants. Insets shows protonema tip Zone, Scale 50 µm. **B)** Colony (top, scale 5 mm) and individual gametophores (bottom, scale 2 mm) from WT and cornichon mutants after four weeks of growth.

Figure S5

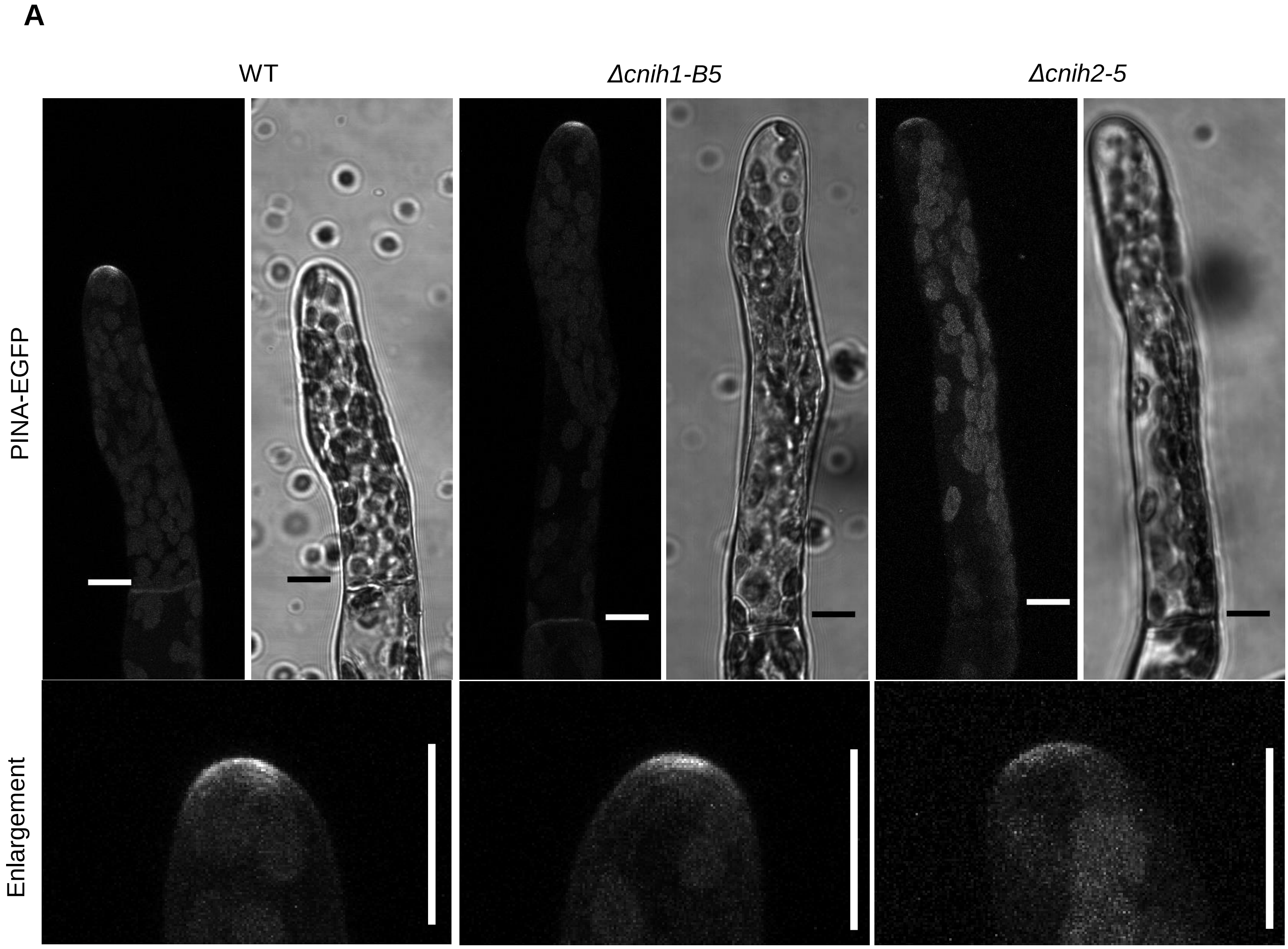


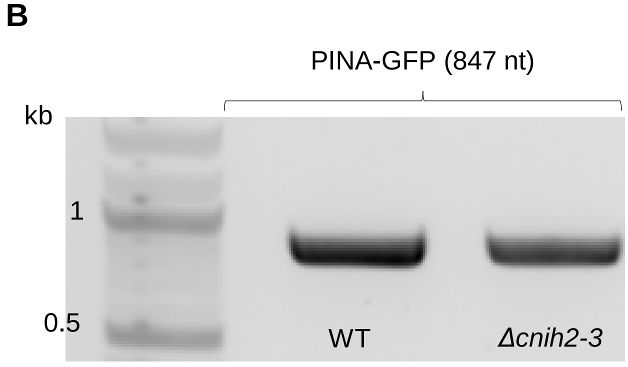


**Figure S5. Subcellular localization of the auxin efflux transporter PINA in other *Δcnih*1 and *Δcnih2* single mutants. A)** Localization of PINA at the tip of the protonema WT apical cell. Fluorescence was maintained in the *Δcnih1-B5* single mutant, but not in the *Δcnih2-5* single mutant. ROI enlargement of PINA-GFP fluorescence at the tip of the apical protonema cells from WT, *Δcnih1-B5*, and *Δcnih2-5* mutant lines. Scale = 10 µm. **B)** PINA-GFP transcript expression in WT and *Δcnih2-3* single mutant line, confirms the presence of the gene.

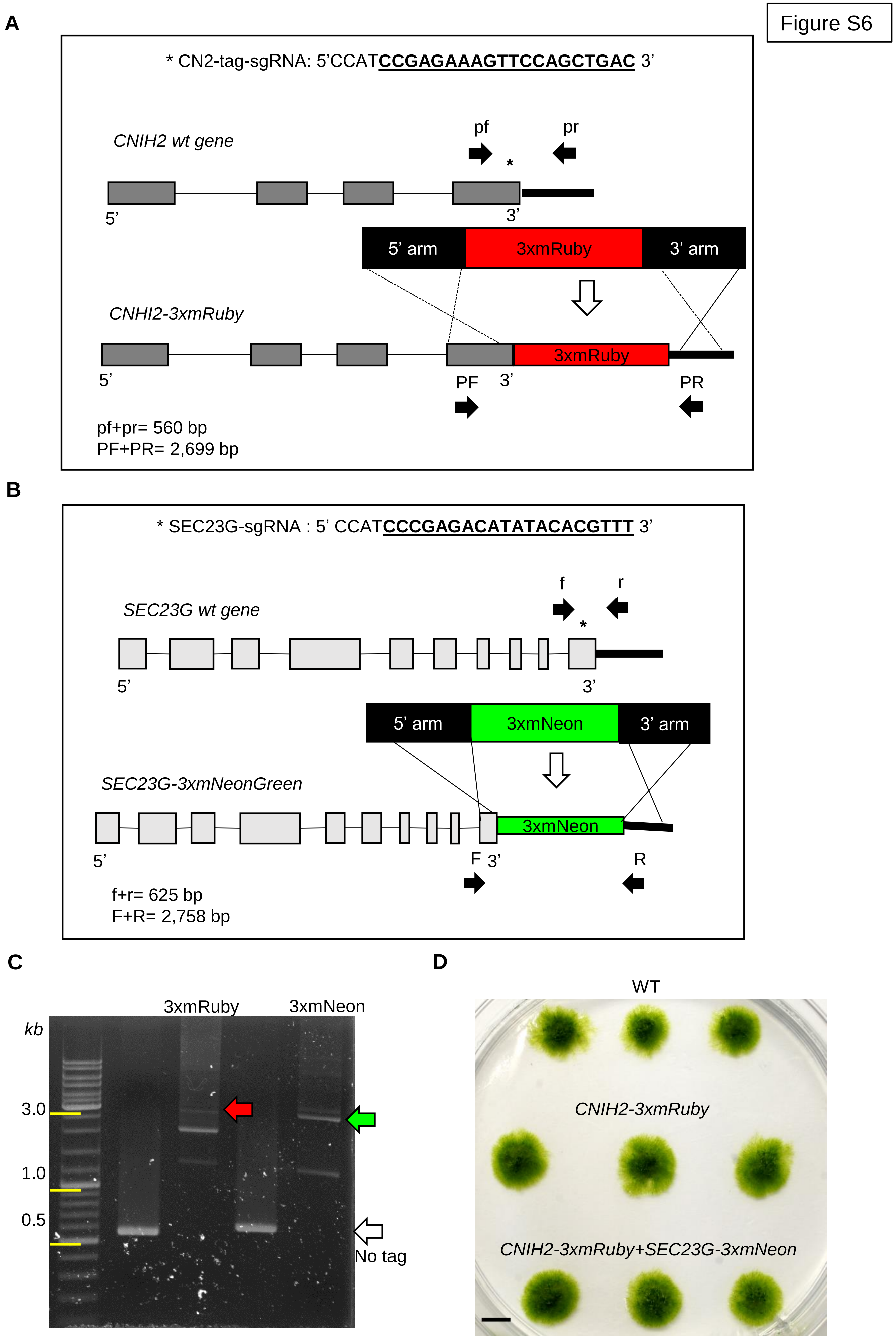
Figure S6
A
* CN2-tag-sgRNA: 5'CCATCCGAGAAAGTTCCAGCTGAC 3'
CNIH2 wt gene
pf
pr
5'
3'
5' arm
3xmRuby
3' arm
CNHI2-3xmRuby
3xmRuby
5'
PF
3'
PR
pf+pr= 560 bp
PF+PR= 2,699 bp
B
* SEC23G-sgRNA : 5' CCATCCCGAGACATATACACGTTT 3'
f
r
SEC23G wt gene
5'
3'
5' arm
3xmNeon
3' arm
SEC23G-3xmNeonGreen
3xmNeon
5'
F
3'
R
f+r= 625 bp
F+R= 2,758 bp
C
3xmRuby
3xmNeon
kb
3.0
1.0
0.5
No tag
D
WT
CNIH2-3xmRuby
CNIH2-3xmRuby+SEC23G-3xmNeon

**Figure S6. Generation of *CNIH2-3xmRuby* and *SEC23G-3xmNeon* Knock-in lines by CRISPR-Cas9 & HDR. A)** Design and representation of *CNIH2-3xmRuby* stable line at the C-terminus of the gene. sgRNA sequence guide for making a double break in DNA (asterisk) and 3xmRuby coding sequence flanked by recombination homolog sequences. Primers pf and pr amplified a fragment of 557 bp by PCR in WT line without inserting the tag. Primers PF and PR amplified a fragment of 2,699 bp by PCR in WT line with the insertion of a 3xmRuby tag. **B)** Design and representation of *SEC23G-3xmNeon* stable line at the C-terminus of the gene. sgRNA sequence guide for making a double break in DNA (asterisk) and 3xmNeon coding sequence flanked by recombination homolog sequences. Primers f and r amplified a fragment of 625 bp by PCR in the WT line without inserting the tag. Primers F and R amplified a fragment of 2,758 bp by PCR in WT line with the insertion of a 3xmNeon tag. **C)** 1% agarose DNA gel showing expected PCR products in the knock-in *CNIH2-3xmRuby* line (red arrow), *SEC23G-3xmNeon* line (green arrow), and without any tag (white arrow). **D)** Images from moss colonies from *CNIH2-3mxRuby* and *CNIH2-3xmRuby+SEC23G-3xmNeon* lines grown together with the WT line. Scale = 2 mm.

Figure S7

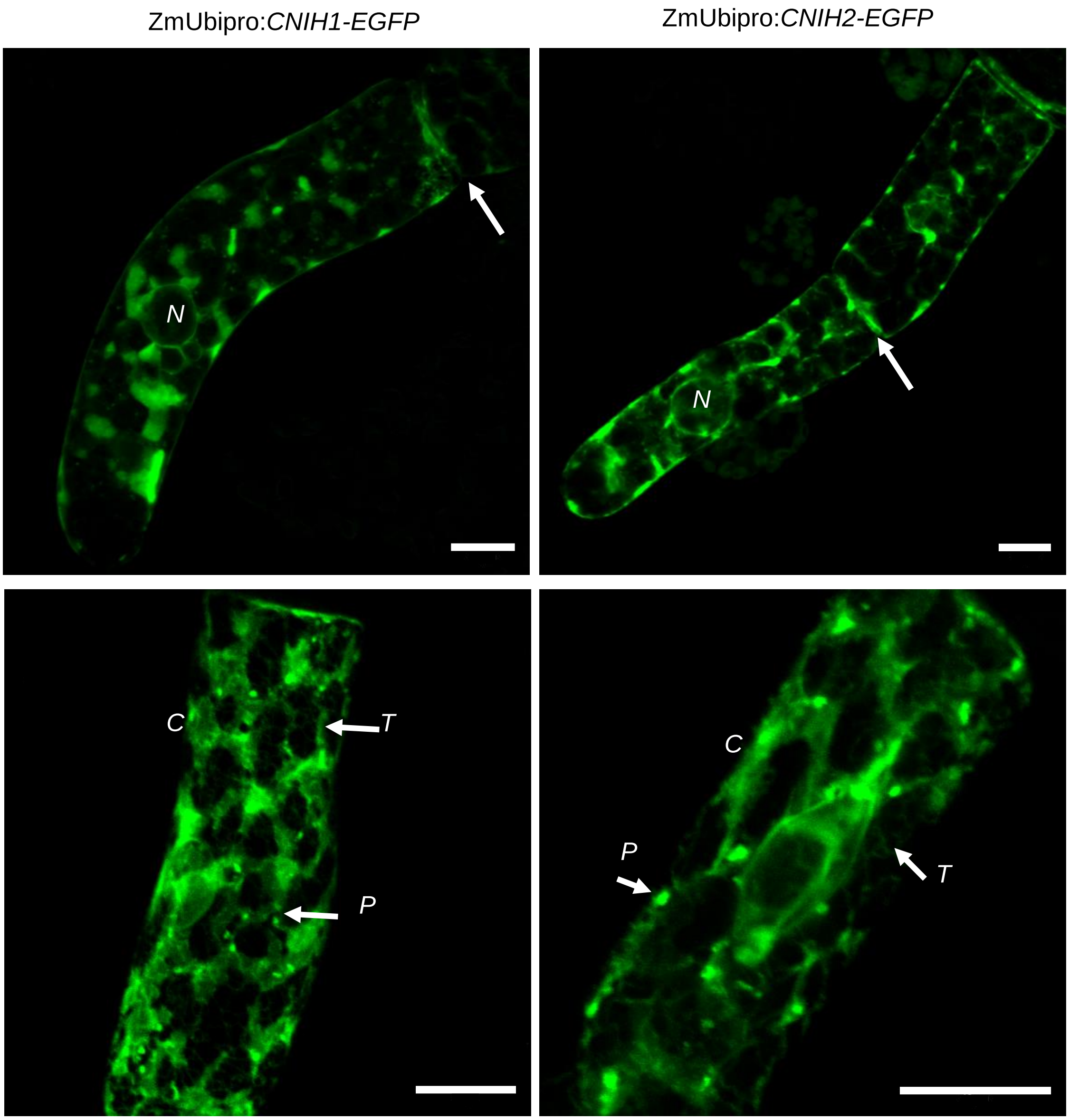


**Figure S7. Overexpression of moss cornichon proteins localized mainly at ER but also in puncta. A)** Confocal images showing the subcellular localization of transiently expressed *ZmUbipro:*CNIH1-EGFP (column left) and *ZmUbipro:*CNIH2-EGFP (column right) in seven-day-old apical protonemal cells. Arrows indicate localization at the cell plate for both proteins (first row). N = nucleus. Identification of moss cornichons at ER subdomains as tubules (T) and cisternae (C), and in puncta below ER (P) (second row), scale 10 µm.

Figure S8

A

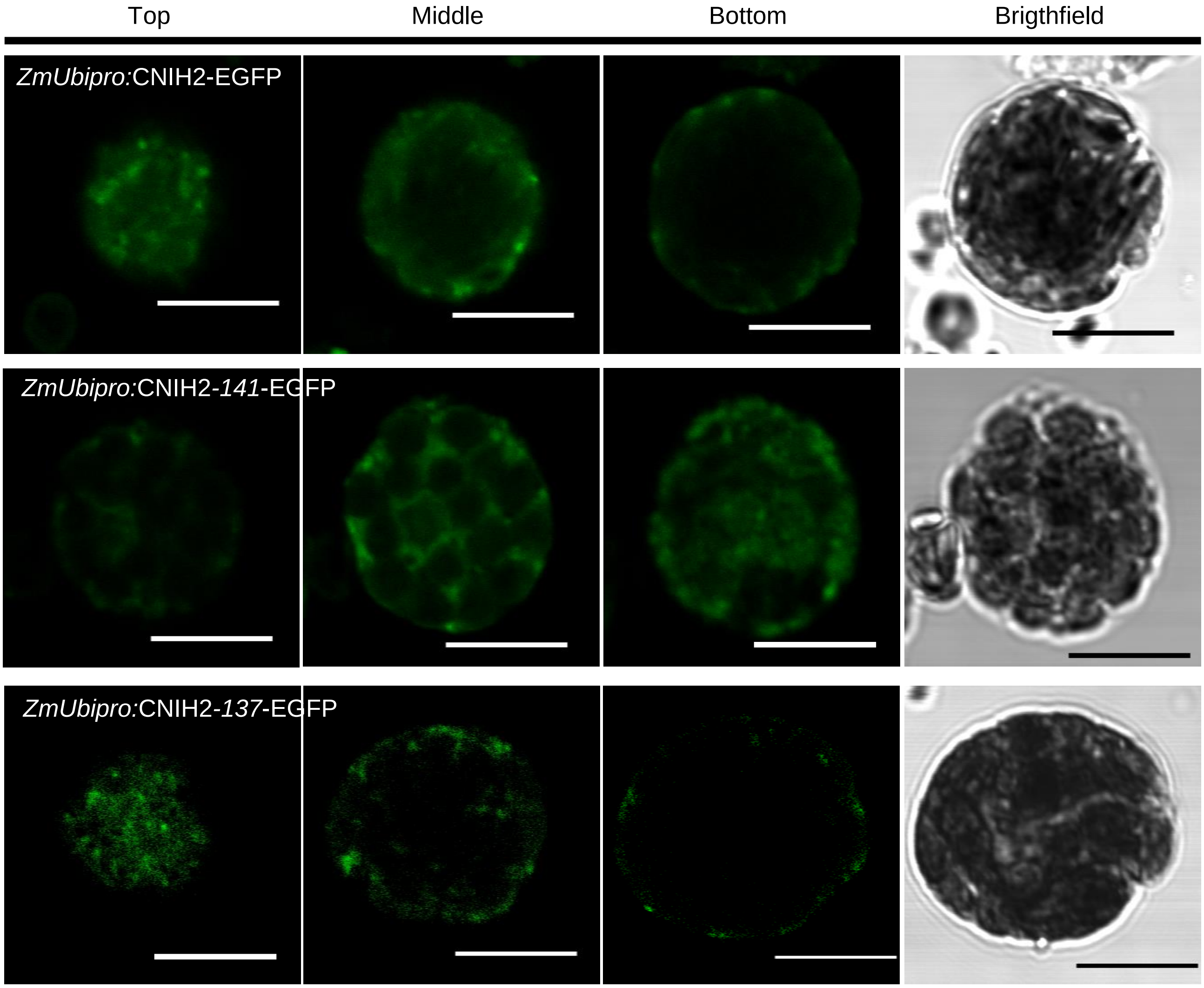
Top
Middle
Bottom
Brigthfield
*ZmUbipro:*CNIH2-EGFP
*ZmUbipro:*CNIH2-*141*-EGFP
*ZmUbipro:*CNIH2-*137*-EGFP

B

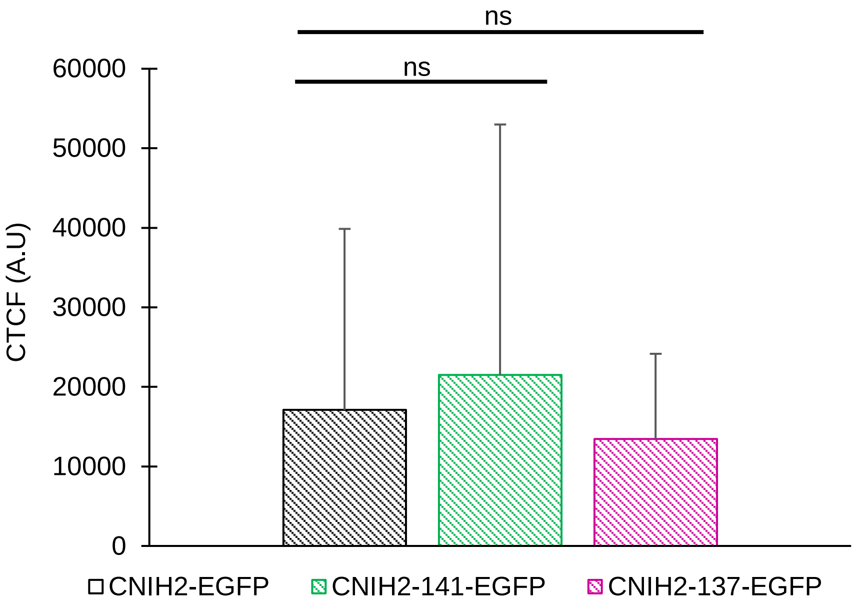
ns
ns
60000
50000
40000
30000
20000
10000
0
CTCF (A.U)
CNIH2-EGFP
CNIH2-141-EGFP
CNIH2-137-EGFP

**Figure S8. Expression of WT CNIH2 and C- terminal truncated proteins in WT moss protoplasts. A)** Confocal images showing expression and localization of transiently overexpression of WT *ZmUbipro:*CNIH2-EGFP (first row) and C-terminus truncated protein versions *ZmUbipro:*CNIH2-141-EGFP (second row) and *ZmUbipro:*CNIH2-137-EGFP (third row) in moss protoplasts at 48 h after transformation, showing no changes in subcellular localization. Images from three individual optical sections (Top, Middle and Bottom) of the protoplasts. Scale 10 µm. **B)** Fluorescence intensity of moss protoplasts transformed with the WT CNIH2 and C-terminus truncated fusion constructs. CTFC = corrected total cell fluorescence. n = 4; data are the mean ± SD t-test was performed for protonema statistics (ns $p \geq 0.05$).

| Table 1. Primers list | | |
|---|---|---|
| Name | SEQUENCE 5' – 3' | References |
| **Cloning** | | |
| attB1 (5' half sequence) | GGGGACAAGTTTGTACAAAAAAGCAGGCT | This study |
| attB2 (5' half sequence) | GGGGACCACTTTGTACAAGAAAGCTGGGT | This study |
| PpCNIH1-For | GTACAAAAAAGCAGGCTTCATGGAGATGGACTTC | This study |
| PpCNIH1-Rev | GTACAAGAAAGCTGGGTCCATGTCTGCGTGGATTG | This study |
| PpCNIH2-For | GTACAAAAAAGCAGGCTTCATGGCTTCCGATCTCC | This study |
| PpCNIH2-Rev | GTACAAGAAAGCTGGGTCCATGTTTGCGTGGATC | This study |
| PpCNIH2-141-Rev | GTACAAGAAAGCTGGGTCTTCCTCATGCTCAAGAATTAAG | This study |
| PpCNIH2-137-Rev | GTACAAGAAAGCTGGGTCAAGAATTAAGTAGACGGCTGC | This study |
| ***PpCNIH1*** **disruption by CRISPR-Cas9 system** | | |
| sgRNACN1-1E-null-Rv | AAACAGCAGCGAGACAACAGCGAA | |
| sgRNACN1-1E-null-Fwd | CCATTTCGCTGTTGTCTCGCTGCT | |
| sgRNACN1-2E-Fwd | CCATTAAGAAGGTAGGTGCAGCCC | This study |
| sgRNACN1-2E-Rv | AAACGGGCTGCACCTACCTTCTTA | This study |
| sgRNACN1-3E-Fwd | CCATACGGAGATCTTCAGTCACCT | This study |
| sgRNACN1-3E-Rv | AAACAGGTGACTGAAGATCTCCGT | This study |
| sgRNACN1 4E-Fwd | CCATCCTCATGCTCCAGAACTAGG | This study |
| sgRNACN1-4E-Rv | AAACCCTAGTTCTGGAGCATGAGG | This study |
| **Genotyping** ***PpCNIH1*** **CRISPR-Cas9 mutants** | | |
| CN1-null-Fwd | CCATTTCGCTGTTGTCTCGCTGCT | This study |
| D-cni1-KO-Rv | GGACGTATGGACTGAATCC | This study |
| ***PpCNIH2*** **Knock-out disruption by Homologous recombination** | | |
| ATTB1-CNI2-M-Fw | GGGGACAAGTTTGTACAAAAAAGCAGGCTGGGTTTAAACGATAGTGAG AGTGAGATGATTGAGG | This study |
| ATTB4-CNI2-M-Rv | GGGGACAACTTTGTATAGAAAAGTTGGGTGGCTCCCGGCTTTCGCTGCT CCTCTC | This study |
| ATTB3-CNI2-4R-Fw | GGGGACAACTTTGTATAATAAAGTTGTAATTCTCTTTGGTTCGTAGCCCA TTGG | This study |
| ATTB2-CNI2-4R-Rv | GGGGACCACTTTGTACAAGAAAGCTGGGTAGTTTAAACAATTCATCTTC GCTTGAACTAC | This study |
| **Genotyping** ***PpCNIH2*** **knock-outs** | | |
| Higro-F | GTCTGTCGAGAAGTTTCTGATCG | |
| Higro-R | CGTCGGTTTCCACTATCGG | |
| CN2-outer-up-F | GGAACGTACATGAGATGTGTCAAG | This study |
| CN2-outer-DW-R | CTCCCTCGTGTACTCCTTCC | This study |
| C2-F | GCTGTTCTTGCTCAACGTTCC | This study |
| C2-R | CTGACTGGACAGATTCCTCATGC | This study |
| Ubi10-F | ACTACCCTGAAGTTGTATAGTTCGG | |
| UbI10-R | CAAGTCACATTACTTCGCTGTCTAG | |
| ***PpCNIH2*** **Knock-in generation by CRISPR-Cas9&HDR** | | |
| CN2-tag-Fwd | CCATCCGAGAAAGTTCCAGCTGAC | This study |
| CN2-tag-Rv | AAACGTCAGCTGGAACTTTCTCGG | This study |
| pENT-CN2-Up-mut-Fw | GAATCTGTaCAGTCAGCTGGAAC | This study |
| pENT-CN2-Up-mut-Rv | GCTGACTGtACAGATTCCTC | This study |
| B1-CNIH2-tag-Fwd | GGGGACAAGTTTGTACAAAAAAGCAGGCTCGTCCTTGCGTCTATCACAT CACG | This study |
| B4-CNIH2-tag-Rv | GGGGACAACTTTGTATAGAAAAGTTGGGTGCATGTTTGCGTGGATCCCC GAGAAAG | This study |
| B3-CNIH2-tag-Fwd | GGGGACAACTTTGTATAATAAAGTTGCGTCCTTGTGACTGTCACACTGA ACC | This study |
| B2-CNIH2-tag-Rv | GGGGACCACTTTGTACAAGAAAGCTGGGTAGCAAGACATGAGCTAGAT ACCAAC | This study |
| **PpCNIH2-3xmRuby line screening** | | |
| cn2-downarm-Fwd | CATGTATCGTTTCTGTCATG | This study |
| cn2-uparm-Rv | GATGTCATGTCAATACCAATG | This study |
| **PpSEC23G-3xmNeon line screening** | | |
| S23g-int-F | GCTACTGATCAATGTTGACTGG | This study |
| S23g-int-R | GAACTAGTTCACTACTCCACG | This study |
| **Transcript expression of PpPINA-EGFP knock-in line** | | |
| EGFP-Fwd | TAAACGGCCACAAGTTCAGCG | This study |
| pinA-Rv | GAGAGGTGCCACCTATTTGCAACC | This study |
